# DNA nanostructure-templated antibody complexes provide insights into the geometric requirements of human complement cascade activation

**DOI:** 10.1101/2023.10.20.562751

**Authors:** Leoni Abendstein, Willem E. M. Noteborn, Luc S. Veenman, Douwe J. Dijkstra, Fleur S. van de Bovenkamp, Leendert A. Trouw, Thomas H. Sharp

## Abstract

The classical complement pathway is activated by antigen-bound IgG antibodies. Monomeric IgG must oligomerize to activate complement via the hexameric C1q complex, and hexamerizing mutants of IgG appear as promising therapeutic candidates. However, structural data have shown that it is not necessary to bind all six C1q arms to initiate complement, revealing a symmetry mismatch between C1 and the hexameric IgG complex, which has not been adequately explained. Here we use DNA nanotechnology to produce specific nanostructures to template antigens, and thereby control IgG valency. These DNA nano-templated IgG complexes can activate complement on cell-mimetic lipid membranes, which enabled us to determine the effect of IgG valency on complement activation without the requirement to mutate antibodies. We investigated this using biophysical assays together with 3D cryo-electron tomography. Our data revealed that the cleavage of complement component C4 by the C1 complex is proportional to the number of antigens. Increased IgG valency also translated to better terminal pathway activation and membrane attack complex formation. Together, these data provide insights into how nanopatterning antigen-antibody complexes influence the activation of the C1 complex and suggest routes to modulate complement activation by antibody engineering. Furthermore, to our knowledge this is the first time DNA nanotechnology has been used to study the activation of the complement system.

## Introduction

Antibodies (Abs) are important effector molecules of the human immune system. Upon binding to antigens, Abs initiate numerous immune responses, ranging from antibody-dependent cell-mediated cytotoxicity, antibody-dependent cellular phagocytosis, to activation of the classical complement pathway.^1-4^ Complement forms part of the humoral innate immune response, and is involved with immune defense against invading pathogens, as well as clearance of cellular debris.^4, 5^ Activation of the classical pathway of complement occurs when the first complement component, the C1 complex, binds to antigen-bound Abs (**Figure 1a**). The C1 complex comprises C1q, which has the ability to bind Abs via the globular head domain (gC1q), and two serine proteases, C1r and C1s, that form a C1r_2_s_2_ heterotetramer (**Figure 1a**).^6^ Binding of C1q to Abs causes activation of C1r and C1s, which proceeds to cleave soluble C4 to form C4b, an opsonin that covalently associates with nearby molecules and membranes.^7, 8^ Some C4b is bound by C2, which is then also cleaved by C1s to form the C4b2b (formerly C4b2a) complex.^9, 10^ C4b2b is a C3 convertase, an enzyme complex that enables complement cascade progression by cleaving C3, resulting in coverage of membranes with multiple covalently-linked opsonins, the release of anaphylatoxins, and formation of the terminal-pathway membrane attack complex (MAC), a pore that perforates the cell membrane.^4-6^

**Figure 1.**
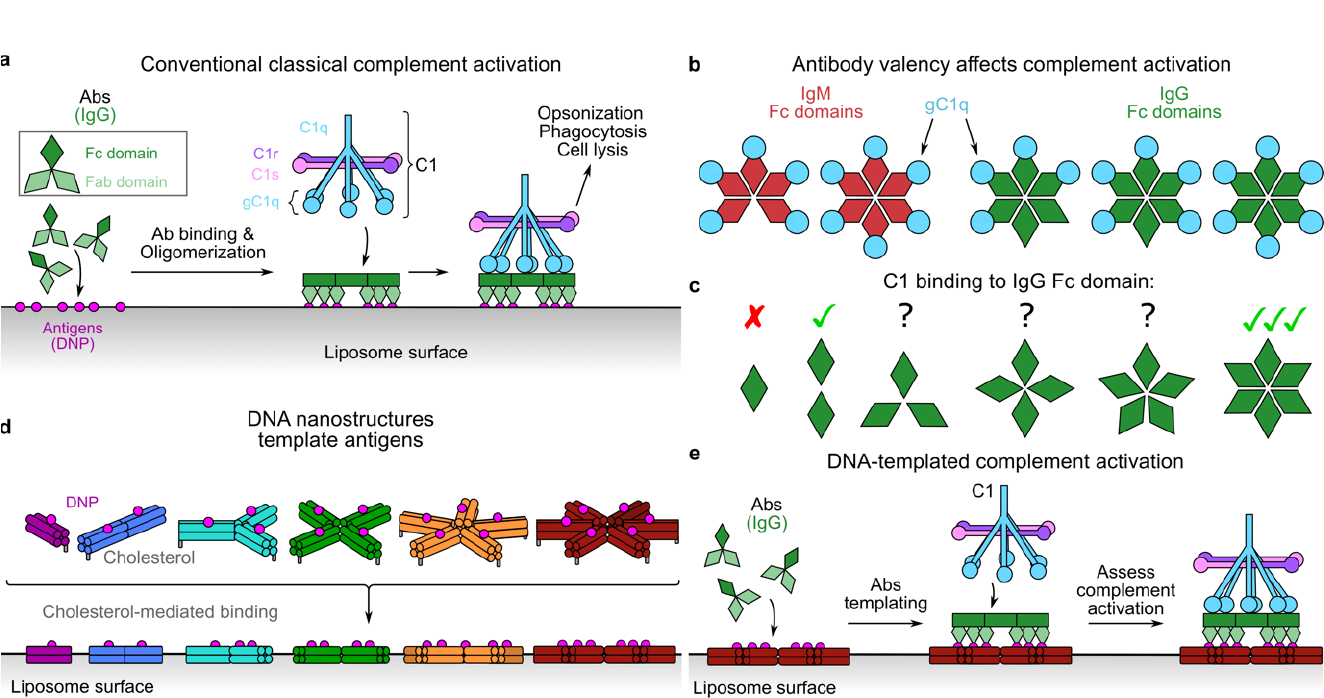
Overview of complement activation and the use of DNA nanostructures to control Ab valency. **a)** Overview of classical complement initiation by the Ab-bound C1 complex. Upon binding to antigens (pink) via Fab domains (light green), the Fc domains of IgG (dark green kites) form nanopatterned platforms (dark green rectangles) that activate the C1 complex. **b**) The Fc domains of IgM (red kites) and IgG (green kites) oligomerize and bind to different numbers of gC1q domains (blue circles). **c**) Valency is known to impact complement initiation, but the geometric requirements of how the IgG Fc monomers (green kites) affect C1 binding, and activation are unknown. **d**) DNA nanostructures can be used to template antigens on lipid bilayers via cholesterol binding. **e**) IgG Abs bind to DNA nanostructure-templated antigens and form platforms of defined valency to explore complement activation.

The classical pathway can be activated by antigen-bound immunoglobulin G (IgG) or IgM Abs. IgM circulates in human serum as a pre-formed pentamer or hexamer,^11^ but IgG exists as a monomer and oligomerization of IgG Abs has been shown to be a requisite for complement activation, with two or more Abs required for C1 binding (**Figure 1b**).^12-14^ More recently, it has been shown that IgG Abs oligomerize via non-covalent interaction between the fragment crystallizable (Fc) regions,^15, 16^ and this has been exploited to develop mutants of IgG subclass 1 with enhanced oligomerization potential that are able to activate complement more effectively than wild-type IgG.^17, 18^ This was achieved by forming hexameric Fc-platforms, to which the C1 complex binds. C1q is a disulfide-linked complex with six flexible arms that bind to the hexameric Fc platform. There is therefore a symmetry match between the Ab-binding C1q complex and the Fc platform. However, structural data have shown that not all C1q arms bind to the hexameric Ab complex,^15, 19^ with four or five arms frequently bound instead of all six (**Figure 1b**). Furthermore, IgM exists preferentially as a pentamer,^11, 20^ but both pentameric and hexameric IgM can bind and activate C1.^7^ These observations reveal a symmetry mismatch between C1 binding and the activating Ab complex, which has not been adequately explained.

Formation of these hexameric platforms is reliant upon the ability of the IgG monomers to bind to antigens at a spacing that provides the means to then form Fc-mediated complexes. This in turn depends on the antigen concentration and proximity, as well as membrane fluidity, epitope location, and IgG subclass. These variables have made it difficult to directly determine the effect of Ab valency on complement activation (**Figure 1c**).

To explore how Ab binding influences complement activation, we use DNA nanotechnology to control antigen valency (**Figure 1d**). DNA can be designed to self-assemble into nanoscale shapes that are chemically accessible, which allows nanometer control over the spacing of functional groups.^21-23^ DNA nanostructures have been previously used to template antigens to determine the effect that spacing and valency have on bivalent Ab binding efficiency with respect to Ab affinities.^24^ However, it has not been used to induce, measure, or control complement activation.

Here, we use DNA nanostructures to pattern antigens with defined valencies that enable us to template Ab complexes and assess the effect of valency on complement activation (**Figure 1d**,**e**). Biophysical characterization of Ab binding and complement initiation revealed that pre-formed Ab complexes activated complement to a greater extent than the same number of Abs left un-patterned. Applying these to liposomal cell mimetics allowed us to assess the degree of membrane rupture caused by complement-mediated MAC pore formation. Furthermore, we augmented these biophysical assays with cryoelectron tomography visualization of DNA nanostructure-mediated complement activation on lipid bilayers in non-purified human serum. Together, these data provide insights into how the C1 complex is activated by nanopatterned antigen-Ab complexes and suggest routes to modulate complement activation by Ab engineering. Furthermore, this research highlights several important caveats that must be taken into consideration for bio-functional DNA nanostructure design before they can be used to modulate extracellular immune system pathways.

## Results

### DNA nanostructure design and characterization

DNA nanostructures were designed to display antigens at specific locations. We used the double-decker tile (DDT) lattice design from Majumder *et al*. as inspiration,^25^ which comprises rigid arms formed from four interconnected double-stranded DNA helices placed on a 2 × 2 grid held together at a central hub (**Figures 2a & S1a**). These arms are symmetric, which helps increase the folding yield.^26^ The DDT design is low-profile, with only ~4 nm height contributed by the two stacked DNA double helices, and the arms are relatively flexible in-plane, which may aid the formation of Ab complexes. The central hub of the design was modified to generate individual tiles containing one (DDT1), two (DDT2), three (DDT3), four (DDT4), five (DDT5) or six (DDT6) arms each comprised of eight different ssDNA strands (S_1-8_; **Figures 2b, c & Table S1**), with two “core” strands that template the number of arms (S_1_ and S_2_), and six “staple” strands, which form the double-decker arms (S_3-8_). The sticky ends of each arm, present in the lattice-forming DDT design,^25^ were removed to inhibit lattice formation (strands labelled “blunt”; **Table S1**). Hinges connecting each arm to a central hub comprised various lengths of unpaired thymine (T) bases at the bending point of the core strands S_1_ and S_2_, which were specific for each design and required by the DNA geometry.^27^ For DDT1 and DDT2 (no bend), no extra Ts were added, whereas three, four, five and seven Ts were added for DDT3 (120° bend), DDT4 (90° bend), DDT5 (72° bend) and DDT6 (60° bend), respectively. To generate DDT1, strands S_3_ and S_4_ were redesigned to form intra-arm base pairs, instead of mediating inter-arm binding, which required the addition of 9 Ts to these strands to allow the 180° bend required (**Figure 2c & Table S1**).

**Figure 2.**
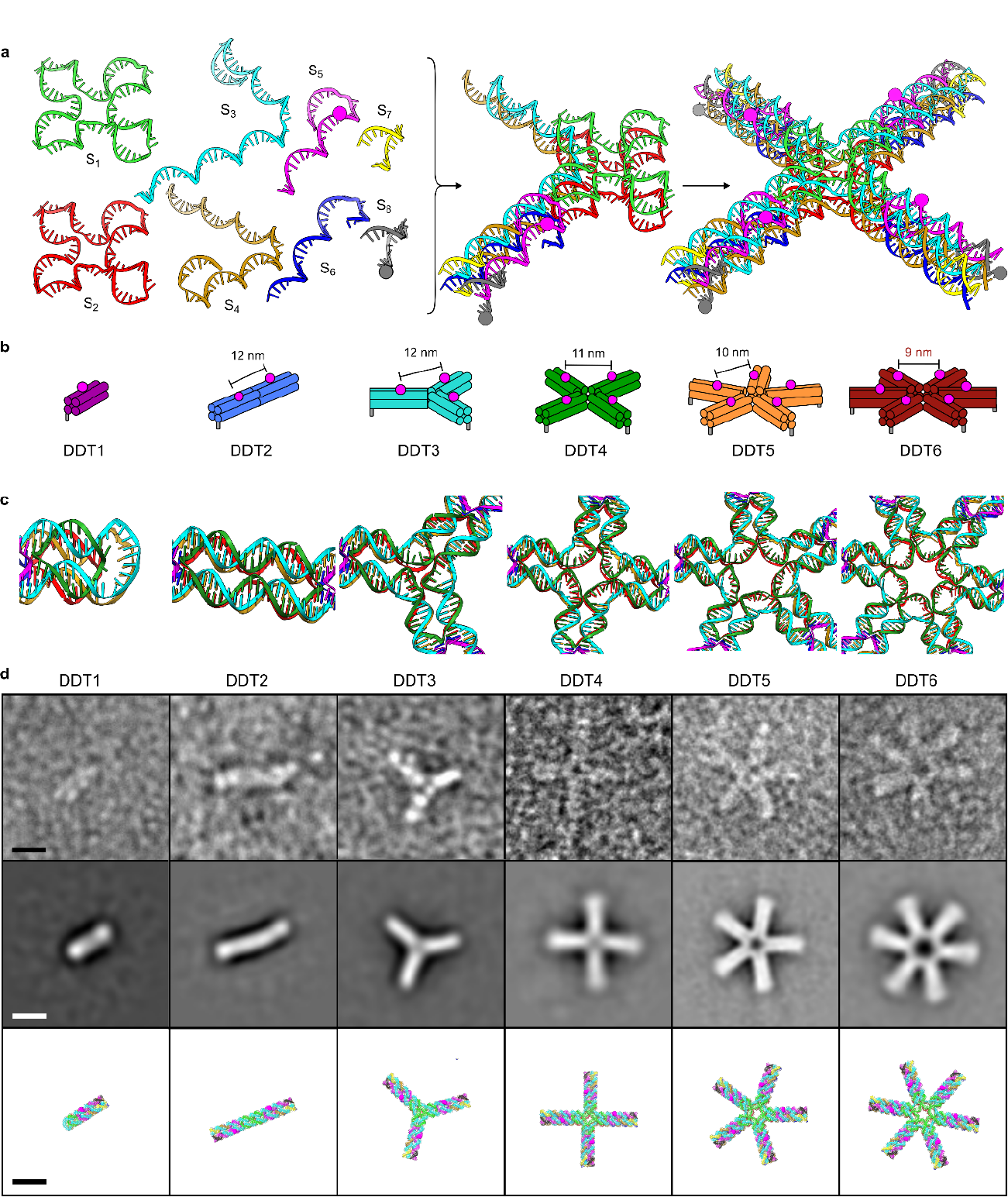
Characterization of DNA nanostructures DDT1-6. **a)** Model of strands S_1-8_ forming DDT4. One of each core strand (S_1_ and S_2_) combine with four copies of staple strands S_3-8_. Antigen positioning on S_5_ is indicated by purple spheres, and cholesterol location on S_8_ is indicated by grey spheres. **b)** Models of DDT1-6, with inter-antigen distances shown. **c)** Detail of modifications to the hub regions necessary to produce DDT nanostructures with defined numbers of arms. **d)** EM micrographs, class averages and models (top, middle, bottom, resp.) of DDT1-6. Scale bars represent 10 nm.

The antigen, 2,4-dinitrophenyl (DNP), was incorporated within the S_5_ strand during synthesis such that it was exposed on the upper face of the DDT (**Figures S1a**,**b**), known as DNP-DDT1-6 henceforth. The S_5_ strand is present and sequence-invariant in all DDT designs. The antigen was placed 6 nm along each DDT arm, as measured from the middle of the nanostructure, yielding antigen-antigen distances of 12, 12, 11, 10, and 9 nm for DDT2, DDT3, DDT4, DDT5 and DDT6, respectively (**Figure 2b**). To activate complement, only one fragment antigen-binding (Fab) domain of the bivalent Ab complex must bind to antigens in order to display the binding site for C1q.^15^ Here, we use human anti-DNP IgG Abs, which are well known to activate complement.^15, 17, 19, 28^

To bind the DDT1-6 to liposomes, the S_8_ strand on the bottom side of the tiles was modified during synthesis to display a 3’ cholesterol group on a tetra-ethylene glycol (TEG) linker (**Figures S1a**,**c**); DNA nanostructures containing cholesterol are labelled DDT1-6-Chol. A positive control comprised a single ssDNA strand modified with both DNP and cholesterol (**Figure S1d Table S1**). This bifunctional strand was synthesized with a 3’ cholesterol group and a 5’ amine group, which was chemically conjugated to a DNP moiety (see Supplementary Information for details).

DNA nanostructures were folded using thermal annealing, which was monitored using agarose gel electrophoresis and size exclusion chromatography (SEC) (**Figures S1e**,**f**). The gel revealed the presence of clear bands corresponding to the folded structures, with minimal unpaired strands at lower molecular weights. Due to the small size of the antigen compared to the overall size of the DNA nanostructure, there was no apparent shift in gel mobility in the presence of DNP. However, the increasing number of arms in each design resulted in clear shifts in both agarose gel mobility and SEC retention volumes, with the main peak of DDT6 eluting at ~0.95 ml and DDT1 eluting at ~1.25 ml, with the remaining designs eluting in between in the expected order. SEC also confirmed the high purity of the folded nanostructures, with a low proportion of unpaired strands eluting at higher retention volumes (~1.73 ml; **Figure S1f**). Consequently, we used the folded DDT nanostructures without further purification. Next, the correct folding of the DDT designs was assessed using negative stain transmission electron microscopy (EM). DDT1-6 were readily identifiable in both electron micrographs and class averages (**Figures 2d & S2a**). These revealed discrete nanostructures with the correct number of arms, and class averages show the limited in-plane flexibility of the rigid arms (**Figure S2b**).

### Binding antibodies to DNA nanostructures

The DDT nanostructures were designed to be used as platforms to bind Abs with distinct valencies. Monoclonal anti-DNP IgG Abs were incubated with DDT1-6 with and without conjugated DNP. Agarose gel electrophoresis showed the appearance of bands and smears with lower motility only in the presence of DNP (**Figure S3a**), indicating Ab binding to DNP-DDTs. To improve our understanding of Ab binding, we performed SEC on DDT nanostructures ± DNP and ± Abs, which revealed Ab binding to all DDT designs only in the presence of all required components (**Figures 3 & S3b**).

**Figure 3.**
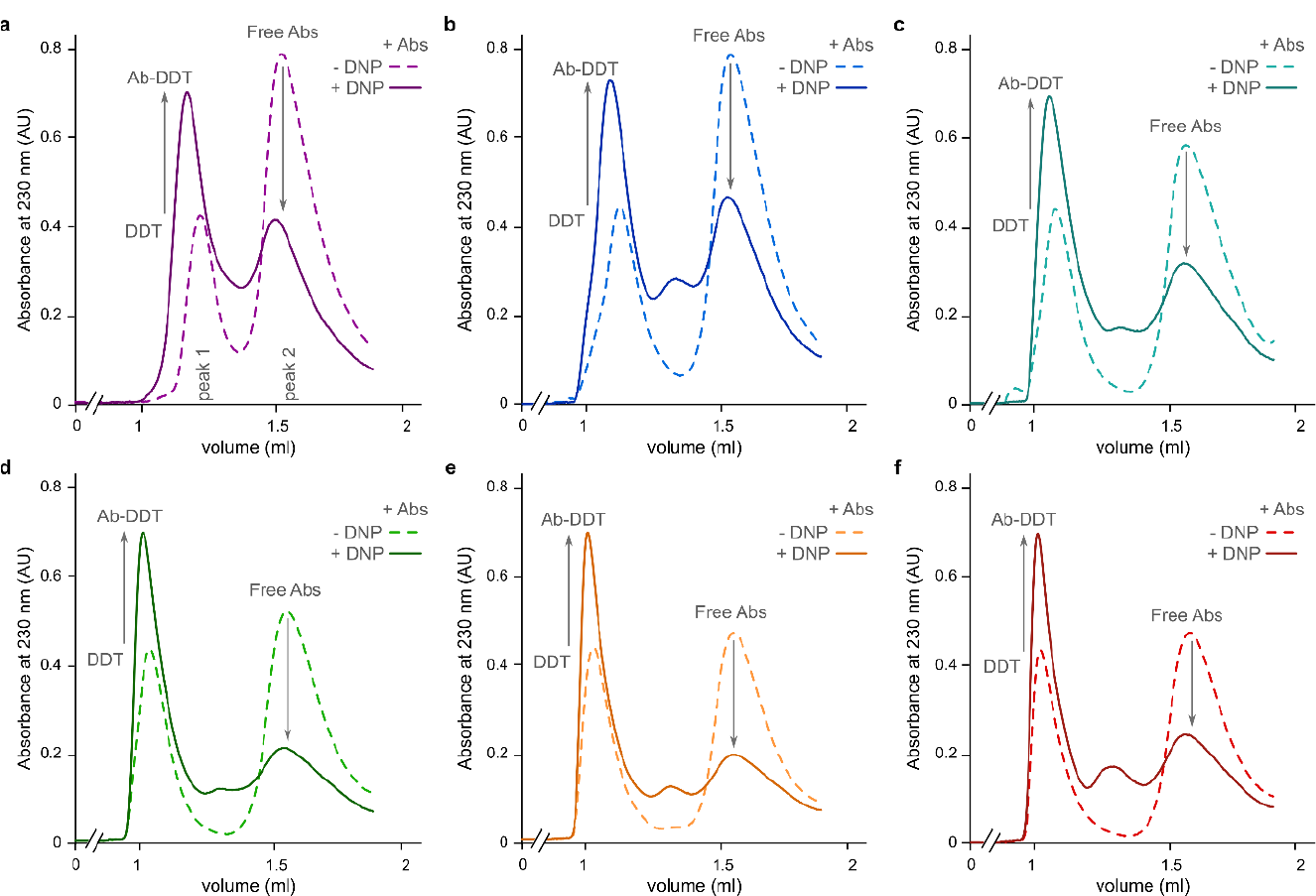
Size exclusion chromatography profiles of DDT nanostructures bound to Abs. **a-f)** Abs added to DDT nanostructures ± DNP antigens for DDT1 **a**), DDT2 **b**), DDT3 **c**), DDT4 **d**), DDT5 **e**) or DDT6 **f**). Arrows indicate changes in peak intensities upon Ab binding.

SEC profiles monitored at 230 nm revealed two prominent peaks, labelled 1 and 2 (**Figure 3a**). In the absence of DNP antigen, peak 1 corresponds to DNA nanostructures (*DDT*), and peak 2 corresponds to free Abs. In the presence of DNP conjugated to the DNA nanostructures, the intensity of peak 1 increases, whilst that of peak 2 decreases, indicating binding of Abs to DNP-DDT designs (*Ab-DDT*; **Figure 3**), and a concomitant reduction in free Abs.

### Pre-patterning antibody complexes using DNA nanostructures enhances complement initiation

To monitor DNA nanostructure mediated C1 binding and activation of the C1r_2_s_2_ serine proteases, we followed complement binding and deposition onto liposome surfaces, which act as cell membrane mimetics. Normal human serum (NHS) was used as a source of complement components. To determine the stability of the DDT nanostructures in the presence of NHS, we incubated DDT6 with either phosphate-buffered saline (PBS) or 10% NHS over a period of 14 h (**Figure 4a**). No difference between the samples in the presence or absence of NHS was apparent, indicating long-term stability of these nanostructures in NHS.

**Figure 4.**
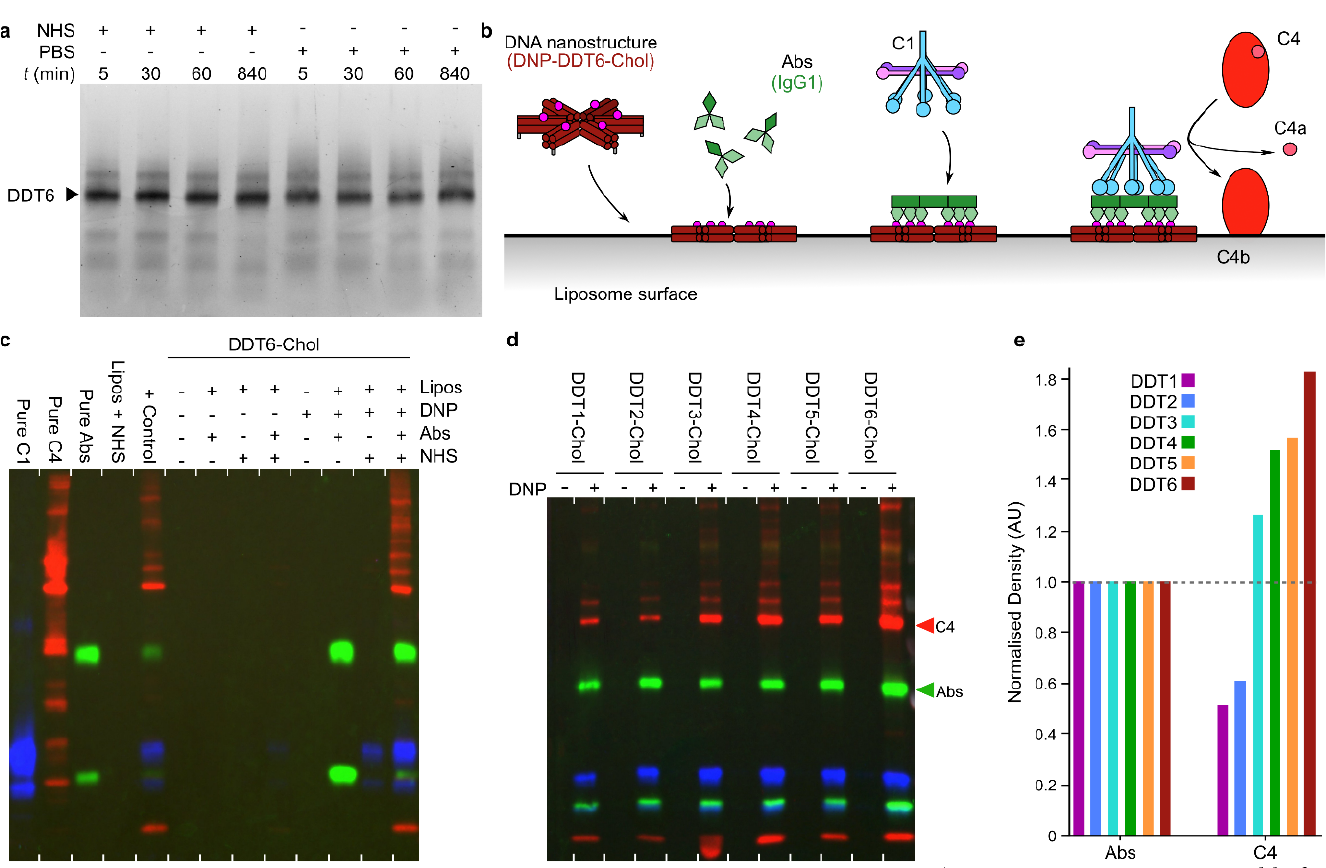
DDT-mediated C1 activation and C4b deposition on liposome membranes. **a)** DDT nanostructures are stable for extended time periods in normal human serum (NHS). DDT6 was incubated with NHS or PBS for times (t) of between 5 min and 14 h. **b)** Schematic showing the initiating steps of the classical complement cascade mediated via DNA nanostructures. At 4°C, the complement cascade is limited to C1 activation and C4b deposition. **c)** Representative western blots overlaid showing the detection of C1q (a component of the C1 complex; blue), C4 (labels both C4 and C4b; red) and IgG Abs (green) on liposomes isolated after incubation with the components shown at the top of the gel. Individual detections on the same blot are shown in **Figure S4b-d**. For the overlaid gels, bands are false-colored for visualization. **d)** Overlaid western blots of C4 co- purified with liposomes after complement activation by DDT1-6-Chol. IgG Abs (green), C1q (blue), and C4 (red), bands are false-colored during overlaying. Blots were treated the same way as **c** and individual detections can be seen in **Figure S4f-h. e)** Quantitative western blot analyses using densitometry analysis of the bands indicated by colored arrowheads in **d**; Ab values were normalized to 1 before C4 values were compared with respect to the normalized Ab values.

DDT6-Chol ± DNP (100 nM final DNP concentration; 16.6 nM DDT6 concentration), and with an excess of anti-DNP IgG Abs (650 nM final concentration) where indicated, were incubated with liposomes and subsequently cooled to 4°C. Ice-cold NHS (10% v/v final concentration), which contains all of the relevant complement components, was then added where indicated. At 4°C, the C1 complex is able to bind to Abs and cleaves C4, but the pathway does not progress beyond C4b deposition (**Figure 4b**).^7, 12^ Liposomes, and any bound material, were then isolated from soluble material by centrifugation.

Silver-stained polyacrylamide gel electrophoresis (PAGE) was used to monitor binding or deposition of proteins and DNA nanostructures onto the liposomes (**Figure S4a**). Gels showed minimal binding of NHS components to liposomes without a source of antigens present (*Lipos + NHS*). This minimal background binding might be caused by incomplete washing of the liposomes after centrifugation. In contrast, extensive protein binding was observed in the presence of the positive control (*+ control*), which comprised a DNA strand modified with both 3’ cholesterol and 5’ DNP. DDT6-Chol was revealed as an isolated band that co-purified with the liposomes, indicating successful binding to the lipid membrane. When incubated with Abs, but without DNP antigens, DDT6-Chol showed minimal background binding of serum proteins. Similarly, as long as Abs were omitted, only a negligible amount of protein bound to the liposome in the presence of NHS, again likely caused by incomplete washing, and compared with positive control the bands are faint. In contrast, DNP-DDT6-Chol (i.e., DDT6 modified with DNP antigens and containing cholesterol) showed co-purification of Abs with the liposomes, indicating DNA-mediated Ab binding to the liposome membrane.

In the presence of DNP-DDT6-Chol, Abs and NHS, extensive protein binding was apparent on the liposome surfaces, indicating successful DNA nanostructure mediated complement activation. Some proteins were also present in samples without Abs but in the presence of DNP-DDT6-Chol and NHS. Here, we also presume this is due to incomplete washing of the liposomes after centrifugation. To verify that we are observing the predicted proteins bound to our liposomes, we performed western blotting for the relevant components; IgG Abs, C1q, and C4 (**Figure 4c**). Each time, the blot was stripped before detection of the next target (**Figure S4b-d**). Clear anti-DNP IgG Abs binding to the positive control and DNP-DDT6-Chol was visible, confirming that the DNA nanostructure was bound to the liposome and able to bind Abs. A small amount of C1q was detected bound to liposomes without Abs on DDT6-Chol ± DNP, but there was no C4 visible, indicating non-activated C1 presumably associating weakly to the DNP-DNA nanostructures or liposomes alone. Overlaying the detection of Abs, C1q, and C4 revealed binding of all three under only two of the conditions; the positive control (*+ control*) and the final lane containing DNP-DDT6-Chol, Abs and NHS with liposomes (**Figure 4c & S4b-d**). This confirms that DDT6 displaying both cholesterol and DNP can activate the complement cascade when bound to the surface of liposomes in the presence of Abs and NHS.

Next, we determined if C4b deposition on liposomes was dependent on the valency of Ab binding. DDT1-6-Chol ± DNP were bound to liposomes before incubation with Abs and NHS at 4°C and purified as described above (**Figures 4d & S4e-h**). Silver-stained gel electrophoresis showed the same result for each DNA nanostructure; proteins bound to the surface of liposomes could only be detected if DDTs contain cholesterol and DNP (**Figure S4e**). We used western blotting to verify that the detected bands were Abs, C1 and C4, which showed copurification of all three components with liposomes for DDT1-6- Chol only in the presence of DNP antigens (**Figure 4d & S4e-h**). Densitometry of the bands was used to determine the effect of Abs valency on C4b deposition. Density values were normalized for Ab concentration in order to directly compare the ability of preformed antigens to activate C1 and stimulate deposition of C4b. This revealed that the amount of C4b deposition was clearly affected by pre-patterning the antibodies into complexes, with a clear correlation of increasing C4b deposition with increasing Ab valency (**Figure 4e**).

### DNA nanostructure-templated antibody complexes activate complement

To enable in situ monitoring of complement activation by DNA nanostructure-mediated Ab complex formation, we bound DDT1-6-Chol to liposome surfaces. Liposomes were formed encapsulating sulforhodamine B (SRB) at a self-quenching concentration.^29^ Complement activation leads to MAC pore formation,^4^ which lyses the liposome and allows dye leakage, resulting in an increase in fluorescence intensity.^7, 30^ DNP-modified DNA nanostructures (DNP-DDT1-6) ± cholesterol were incubated with liposomes in PBS and thereby functioned as the only antigen source for classical complement activation.

Different concentrations of DNP-DDT1-6, ranging from 10 nM to 100 nM total DNP (e.g. for 100 nM total DNP concentration, 100 nM, 50 nM, 33.3 nM, 25 nM, 20 nM and 16.6 nM of DDT1, DDT2, DDT3, DDT4, DDT5 and DDT6, respectively, were used), were incubated with the liposomes for 10 min before an excess of anti-DNP IgG Abs (350 nM final concentration) was added. After a 100 s incubation to allow Abs to bind to the DNA nanostructure-templated DNP antigens, NHS (10 % v/v final concentration) was added, and the fluorescence intensity was monitored. No complement activation occurred if no Abs were added or no antigen was present, and DNA nanostructures did not lyse the liposomes on their own. In contrast, the positive control, comprising a DNA strand modified with both 3’ cholesterol and 5’ DNP, showed clear complement activation (**Figures 5 & S5a**).

**Figure 5.**
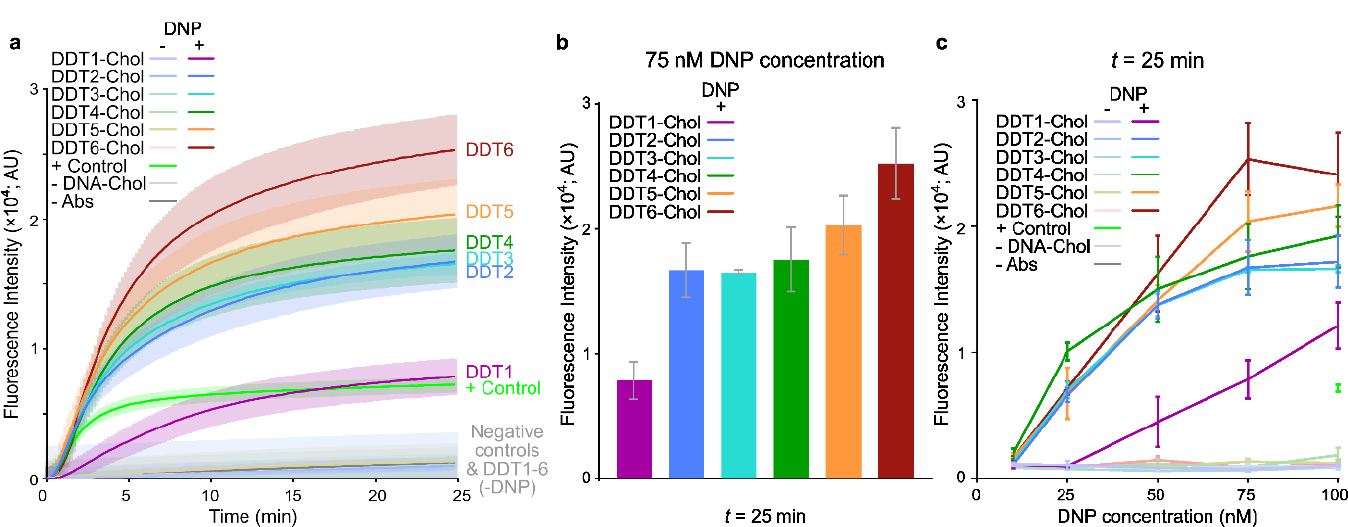
Liposome lysis assays reveal DNA nanostructure-templated Ab complexes can activate complement. **a)** Liposome lysis assay showing an increase in fluorescence caused by DNA nanostructure-templated Ab binding at 75 nM DDT. **b)** Plots of the fluorescence intensity after 25 min of the assay at 75 nM DDT. **c)** Fluorescence intensity of DNP-DDT1-6-Chol mediated lysis, as measured after 25 min. Error bars in all panels represent the standard error of 3 independent replicates.

At low (10 nM) DNP-DDT concentrations, no complement activation occurred for any of the designs (**Figure S5b**). However, above 25 nM total DNP, we observed complement activation mediated by DNA-nanostructures. DNA nanostructure-templated Ab complexes with two or more antigens (DNP-DDT2-6-Chol) were clearly more effective than monomeric DNP-DDT1-Chol at activating complement (**Figures 5c & S5c-f**). With a concentration of 50 nM DNP, the differences between DNP-DDT1-Chol and DNP-DDT2-6-Chol were more apparent, with DNP-DDT2-6-Chol all activating complement equally efficiently (**Figure S5d**). With 75 nM DNP, DNP-DDT6-Chol was the most effective complement activator, followed by DNP-DDT5-Chol, DNP-DDT4-Chol, and then DNP-DDT2-Chol and DNP-DDT3-Chol with approximately equal efficiency (**Figures 5 & S5e**). At the highest concentration of 100 nM DNP, the differences between the nanostructures were reduced, with DNP-DDT1-Chol able to activate complement nearly as efficiently as DNP-DDT2-Chol, although the overall trend of higher oligomeric states resulting in increased complement activation remained (**Figures 5c & S5f**).

### Visualization of DNA nanostructure-mediated complement activation using cryoET

To visualize complement activation by the DNA nanostructure-templated Ab complexes on liposomes, we used cryo-electron tomography (cryoET). Liposomes were incubated with 50 nM DNP-DDT6-Chol (**Figure 6a**). Next, an excess of anti-DNP IgG Abs (final concentration 350 nM) was added and subsequently cooled to 4°C before the addition of ice-cold NHS (1.5% v/v final concentration), to halt the complement cascade at C4b deposition.^7, 12^ Liposome samples were prepared for cryoET that contained (*left to right;* **Figure 6a**,**b**); DNP-DDT6 ± cholesterol, with anti-DNP IgG Abs, or with all the required components cooled to 4°C.

**Figure 6.**
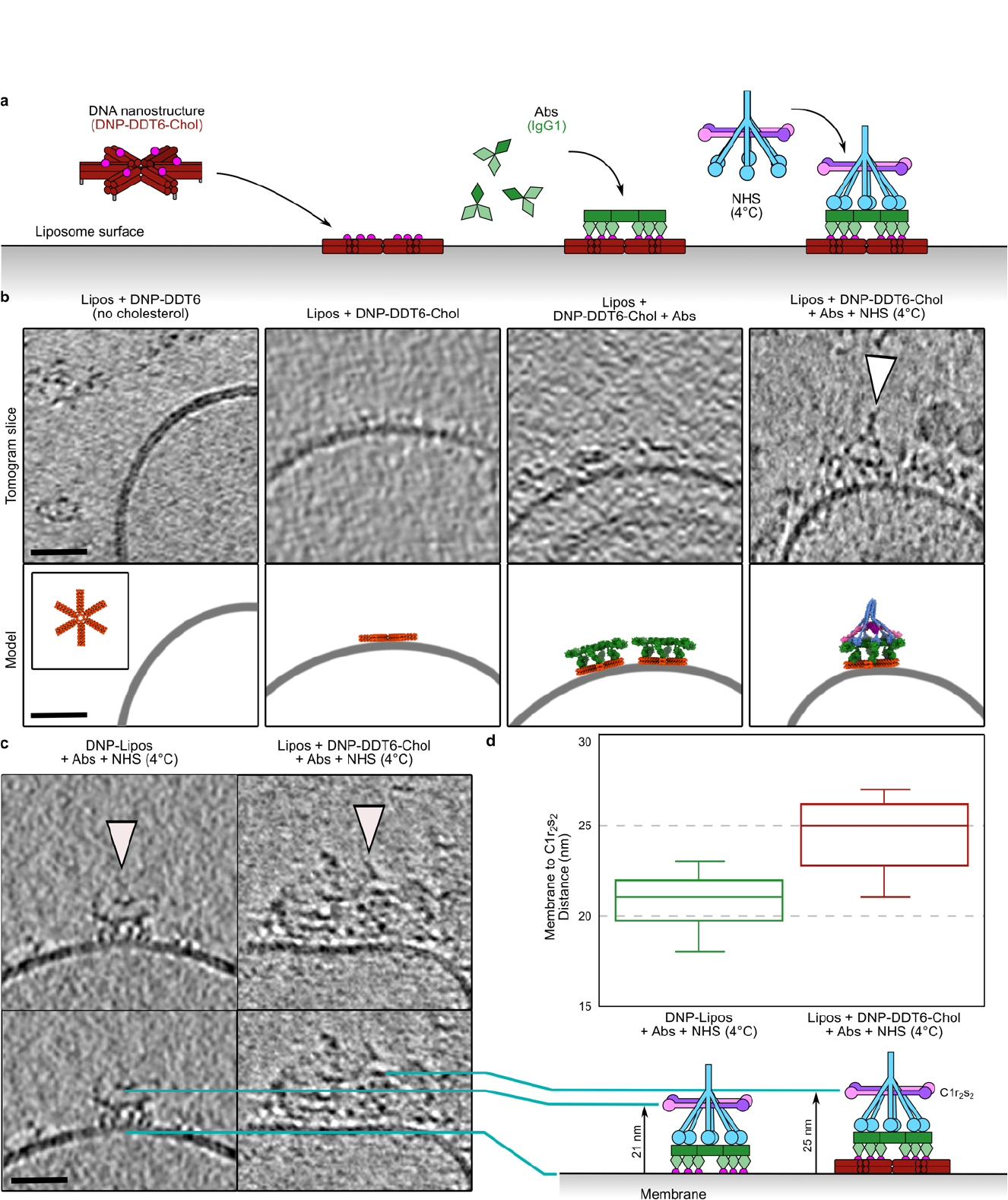
Cryo-electron tomography of DNA nanostructure mediated complement activation. **a)** Schematic of steps required for C1 binding on liposome associated DDT6. **b)** Slices, ~10 nm thick, through cryo-electron tomograms at each step of the schematic shown in **a. c)** Difference in height between C1 bound to Abs on DNP-coated liposomes (left) and DNP-DDT6-Chol nanostructures recruited Abs on liposomes **d)** Box plot of height measurements from the liposome membrane to the C1r_2_s_2_ platform visible in the tomogram slices (n=10 for both). The schematics below show how the height was measured. Scale bars all represent 30 nm.

In the absence of cholesterol, no DNP-DDT6 was observed bound to the liposome membranes, although material with the correct dimensions for solution-phase DNP-DDT6 was visible in the vitrified medium (**Figure 6b**). However, DNP-DDT6- Chol was observed on the liposome surfaces visible as low-profile density. Upon the addition of anti-DNP IgG Abs to liposomes with DNP-DDT6-Chol, clear platforms appeared (**Figures 6b & S6a**) that corresponded closely with IgG platforms previously observed using cryoET.^19, 31^ Finally, after incubation with chilled NHS, C1 complexes were clearly visible on top of Abs platforms recruited by DNA nanostructure-templated Ab complexes (**Figures 6b & S6b**). Interestingly, these C1 complexes appeared farther from the membrane than the IgG-C1 complexes previously visualized when bound to antigenic membranes (**Figures 6c & S6c**).^19^ On liposomes binding anti-DNP IgG Abs directly, the DNP antigens were linked onto the lipid. Measuring the distance from the lipid membrane to the C1r_2_s_2_ protease platform, clearly visible in tomograms as a platform parallel to the liposome membrane (**Figure 6c,d**), yielded different values. Whereas, for C1 bound to Abs on the surface of DNP displaying liposomes and C1 bound to Abs recruited by DNP-DDT6-Chol on the surface of liposomes of 21 and 25 nm, respectively (**Figure 6d**). Consequently, when bound to DNA nanostructure-templated Ab complexes, C1 was 4 nm farther from the membrane, approximately the same height that two stacked DNA double helices will add.

## Discussion

Antibody engineering has led to the development of monomeric IgG Abs with an enhanced propensity to oligomerize.^15^ The design and development of these supramolecular Ab complexes required extensive rational and random mutagenesis,^15, 17^ which led to a single hexameric species held together via non-covalent interactions between the Fc domains.^16^ This so-called Hexabody® technology is being developed as a potential therapeutic to enhance immune responses, such as the activation of the complement system.^17^ However, other oligomeric states exist with a multitude of geometric nanopatterns, and it is not yet clear how many of these are sampled by antibodies, or indeed capable of activating the complement system (**Figure 1**). By using differential combinations of mutations, Strasser et al. could modulate IgG mediated complement activation based on oligomeric state.^28^ These data indicated that a minimum of four C1q binding sites were required for complement activation, in contrast to previous data, which showed that two Abs is sufficient.^12-14^ This difference may be due to the extensive mutations required to induce protein oligomeric state.^28^

By using DNA as a nanopatterning scaffold, we were able to template wild-type (non-mutated) IgG Abs on DNA nanostructures with distinct valencies, and then scan valency space with fewer structural perturbations than Abs engineering requires (**Figure 2**). Our DNA nanostructure design was chosen to be flexible and low-profile to minimize any structural perturbations besides the antigen valency. Previously, antigens have been nanopatterned on rigid DNA nanostructures,^24^ but these, even when placed on an “idealized” hexameric array, were not shown to activate complement. Whether this is due to some inherent flexibility required for C1 activation, or due to the much larger DNA design, is currently not known. Complement activation by the C1 complex is induced upon C1q binding, which leads to the activation of C1r within the heterotetrametric C1r_2_s_2_ platform. It is not yet known if C1r activation proceeds via auto-activation (i.e., C1r cleaves the adjacent C1r within the same C1r_2_s_2_ platform), or cross-activation (C1r cleaves the C1r of an adjacent C1 complex). By minimizing the size of our DNA nanotemplates, we have allowed either to occur and not inhibited cross-activation; indeed, the DNA nanostructures are able to diffuse along the lipid membrane, and this will allow lateral C1 interactions that have been previously posited to be present during C1 activation.^19^ Furthermore, C1 produces multiple copies of C4b, an opsonin that is deposited on the lipid membrane,^7^ which then binds to C2 before cleavage by C1s to form the C4b2b C3 convertase enzyme. This means that C4b must be able to diffuse away from the C1 complex to both function as an opsonin and to allow binding of C2 before cleavage. Here again, the small size of our DDT nanostructures minimizes steric perturbations to this enzymatic reaction. In contrast, larger DNA nanostructures may limit or even inhibit these requirements, leading to stalled complement progression and no MAC pore formation.

Clearly, the impact of the shape and chemistry of DNA nanostructures are important parameters to take into consideration when designing DNA nanotemplates. For use in biological or biophysical assays, the scalability and variability of the DNA design should also be taken into account. The DDTs utilized in this study were designed such that the modifications required to change the antigen valency were minimal. Unlike nanostructures based on DNA origami, DDTs are composed of 8 small single-stranded DNA (ssDNA) oligonucleotides and do not require a ssDNA scaffold. They are also based on a symmetric design,^25, 26^ which greatly reduces the number of strands required. Furthermore, the DNP antigen was incorporated during oligo synthesis specifically at the location required for Ab binding, and this strand was present in all DDT designs. This has two benefits: the financial outlay is reduced, as most of the same strands can be used for each design; and there are minimal differences between the designs (only two of the 8 strands were altered for each design; **Table S1**), which allowed for more robust comparisons between their complement-activating abilities.

Comparing the DDT designs revealed that C4b deposition is proportional to the number of antigenic arms within the DDT designs (**Figure 4**), and MAC pore formation increased concurrently with the number of antigenic arms present within the DDT designs (**Figure 5**), although this was less apparent than for C4b deposition. DDT1 was the worst construct for both C4b deposition (**Figure 4**) and MAC pore formation (**Figure 5**), presumably due to the requirement of IgG Abs to bind to each DDT1 individually, and then rely on lateral diffusion to form higher order Ab oligomers that are capable of C1 binding and activation.^28^ DDT2 was also poor at C4b deposition; indeed, C4b deposition improved with higher valency (**Figure 4e**). However, this trend did not translate to MAC pore formation. Between 25-50 nM, DDT2-6 all performed similarly well at membrane lysis (**Figure 5**). This discrepancy may be caused by a reduced ability of C1 to activate on nanopatterns with lower valency, but still produce sufficient C4b to initiate the complement cascade. This proteolytic cascade results in a positive feedback loop that may mask the differences in C4b deposition by amplifying the cascade such that equivalent numbers of MAC pores are formed, leading to equivalent liposome lysis. From 25 nM DNP and above, DDT2-6 were able to lyse liposomes, whereas DDT1 needed at least 50 nM DNP to achieve lysis. Interestingly, at 25 nM as well as 50 nM, all five designs (DDT2-6) were broadly similar in their membrane lysing abilities, indicating that two proximal antigens are indeed sufficient to display Abs in a conformation able to activate complement, which is in agreement with previous data.^12-14^

Why two antigens are sufficient to activate the classical complement pathway and lyse liposomes is not yet clear. We cannot exclude the possibility that two antibody-bound DNA nanostructures are able to align and enable C1 binding. Indeed, this possibility was included in the design to allow cross-activation of the C1 complexes, as described above. C1 binding to multiple DNA nanostructure-templated Abs complexes is presumably how DNP-DDT1-Chol is able to activate complement above 50 nM concentration. However, below 50 nM, DDT1 is not able to activate complement, indicating that at this concentration the other designs are also sufficiently separated such that only one C1 complex binds per DNA nanostructure. Above 50 nM, DDT6 was superior to DDT5 at liposome lysis, with DDT2-4 achieving similar levels of lysis. Presumably, DDT6-templated hexameric antibody platforms can bind all 6 arms of the C1q complex to achieve maximal C1 activation. Previous models of C1 activation have utilized either a “compaction” of the C1q arms,^19^ or a sliding motion within the C1r_2_s_2_ platform upon C1q binding to antibodies.^7^ To understand the structural biology of DNA nanostructure mediated complement activation we attempted to image the samples using cryo-electron tomography (**Figure 6**). Tomographic volumes revealed the C1 complexes binding to DNA nanostructure-templated antibody platforms, although we were unable to resolve a 3D map of the entire complex. We posit that this was due to the flexibility of the system; previous maps have been hampered by flexibility and therefore only partially resolved,^19^ and we have added a layer of complexity with the addition of flexible DNA nanostructures. Nevertheless, we were able to image stepwise reconstitution of liposome-bound DDT6-chol, Abs binding to DNP-DDT6-chol, and C1 complex association to the DNA nanostructure-templated Ab complexes (**Figure 6**).

Our work represents the first time that DNA nanostructure-templated antibodies have been used to activate the complement system. By varying the valency of the nanotemplates, we could control the geometric parameters of complement activation. Recent work has shown that higher Fc platforms behave differently,^31^ with taller IgG3 complexes better able to activate complement than shorter IgG. The work described herein shows a route to explore this further, by systematically varying the height of the initiating complex via the use of DNA templates of a defined height. This may lead to a greater understanding of how the epitope location relates to complement activation.^18, 32, 33^

We also discovered a correlation between antibody valency and C4b deposition, which implies a potential mechanism to switch complement from an inflammatory response to silent clearance. Although inflammation is generally desired to fight infections, silent clearance of cellular debris is optimal to limit sensitization of the immune system towards benign cellular material, which therefore limits autoimmune responses.^34-36^ By controlling C4b deposition in vivo, it may be possible to limit opsonization to induce phagocytosis without the formation of MAC pores. DNA nanostructures can be targeted to distinct cell types via binding motifs such as aptamers ^37^ or directly displaying ligands for receptor binding,^38^ and are stable in serum and in vivo (**Figure 4**).^39, 40^ The use of DDT2-4 to induce *in vivo* dimerization, trimerization or tetramerization of IgG, instead of hexamerization Abs, would be a novel route to induce sub-lytic complement activation.

## Author Contribution

LA performed all biochemical assays. LA and THS collected cryoET data. LA, WEMN, LSV and THS designed and generated DNA nanostructures. DJD, FSB and LAT constructed vectors, and expressed and purified IgG. LA and THS performed tomogram reconstructions. LA and THS wrote the paper. All authors contributed to paper revisions. THS supervised the study.

## Notes

The authors declare no competing interests.

## Acknowledgements

This research was supported by the following grants to THS: European Research Council Grant 759517; The Netherlands Organization for Scientific Research Grants VI. Vidi.193.014. This work benefited from access to the Netherlands Centre for Electron Nanoscopy (NeCEN) at Leiden University, an Instruct-ERIC center with assistance from Christoph A. Diebolder.

## References

(1) de Taeye, S. W.; Bentlage, A. E. H.; Mebius, M. M.; Meesters, J. I.; Lissenberg-Thunnissen, S.; Falck, D.; Senard, T.; Salehi, N.; Wuhrer, M.; Schuurman, J.; et al. FcgammaR Binding and ADCC Activity of Human IgG Allotypes. Frontiers in Immunology 2020, 11, 740. DOI: 10.3389/fimmu.2020.00740

(2) Tay, M. Z.; Wiehe, K.; Pollara, J. Antibody-Dependent Cellular Phagocytosis in Antiviral Immune Responses. Frontiers in Immunology 2019, 10, 332. DOI: 10.3389/fimmu.2019.00332

(3) van Erp, E. A.; Luytjes, W.; Ferwerda, G.; van Kasteren, P. B. Fc-Mediated Antibody Effector Functions During Respiratory Syncytial Virus Infection and Disease. Frontiers in Immunology 2019, 10, 548. DOI: 10.3389/fimmu.2019.00548

(4) Ricklin, D.; Hajishengallis, G.; Yang, K.; Lambris, J. D. Complement: a key system for immune surveillance and homeostasis. Nature Immunology 2010, 11 (9), 785–797. DOI: 10.1038/ni.1923

(5) Merle, N. S.; Church, S. E.; Fremeaux-Bacchi, V.; Roumenina, L. T. Complement System Part I - Molecular Mechanisms of Activation and Regulation. Frontiers in Immunology 2015, 6, 262. DOI: 10.3389/fimmu.2015.00262

(6) Zwarthoff, S. A.; Ruyken, M.; Magnoni, S.; Vidarsson, G.; Aerts, P. C.; de Haas, C. J.; Rooijakkers, S. H. M. Defining the molecular interplay between antibodies and complement in bacterial infections. Molecular Immunology 2017, 89, 154–155. DOI: 10.1016/j.molimm.2017.06.113

(7) Sharp, T. H.; Boyle, A. L.; Diebolder, C. A.; Kros, A.; Koster, A. J.; Gros, P. Insights into IgM-mediated complement activation based on in situ structures of IgM-C1-C4b. Proceedings of the National Academy of Sciences of the United States of America 2019, 116 (24), 11900–11905. DOI: 10.1073/pnas.1901841116

(8) Mortensen, S.; Kidmose, R. T.; Petersen, S. V.; Szilagyi, A.; Prohaszka, Z.; Andersen, G. R. Structural Basis for the Function of Complement Component C4 within the Classical and Lectin Pathways of Complement. The Journal of Immunology 2015, 194 (11), 5488–5496. DOI: 10.4049/jimmunol.1500087

(9) Bohlson, S. S.; Garred, P.; Kemper, C.; Tenner, A. J. Complement Nomenclature-Deconvoluted. Frontiers in Immunology 2019, 10 (1308), eCollection 2019. DOI: 10.3389/fimmu.2019.01308

(10) Krishnan, V.; Xu, Y.; Macon, K.; Volanakis, J. E.; Narayana, S. V. The structure of C2b, a fragment of complement component C2 produced during C3 convertase formation. Acta Crystallographica Section D 2009, 65 (Pt 3), 266–274. DOI: 10.1107/S0907444909000389

(11) Hughey, C. T.; Brewer, J. W.; Colosia, A. D.; Rosse, W. F.; Corley, R. B. Production of IgM Hexamers by Normal and Autoimmune B Cells: Implications for the Physiologic Role of Hexameric IgM. The Journal of Immunology 1998, 161 (8), 4091–4097. DOI: 10.4049/jimmunol.161.8.4091

(12) Borsos, T.; Rapp, H. J. Complement Fixation on Ceil Surfaces by 19S and 7S Antibodies. Science 1965, 150, 505–506.

(13) Rosse, W. F. The fixation of C1 by autoimmune antibody and heterologous anti-IgG antibody. The Journal of Clinical Investigation 1971, 50, 772–733.

(14) Doekes, G.; Van Es, L. A.; Daha, M. R. Influence of aggregate size on the binding and activation of the first component of human complement by soluble IgG aggregates. Immunology 1982, 45 (705-713).

(15) Diebolder, C. A.; Beurskens, F. J.; de Jong, R. N.; Koning, R. I.; Strumane, K.; Lindorfer, M. A.; Voorhorst, M.; Ugurlar, D.; Rosati, S.; Heck, A. J.; et al. Complement is activated by IgG hexamers assembled at the cell surface. Science 2014, 343 (6176), 1260–1263. DOI: 10.1126/science.1248943

(16) Strasser, J.; de Jong, R. N.; Beurskens, F. J.; Schuurman, J.; Parren, P.; Hinterdorfer, P.; Preiner, J. Weak Fragment Crystallizable (Fc) Domain Interactions Drive the Dynamic Assembly of IgG Oligomers upon Antigen Recognition. ACS Nano 2020, 14 (3), 2739–2750. DOI: 10.1021/acsnano.9b08347

(17) de Jong, R. N.; Beurskens, F. J.; Verploegen, S.; Strumane, K.; van Kampen, M. D.; Voorhorst, M.; Horstman, W.; Engelberts, P. J.; Oostindie, S. C.; Wang, G.; et al. A Novel Platform for the Potentiation of Therapeutic Antibodies Based on Antigen-Dependent Formation of IgG Hexamers at the Cell Surface. PLoS Biology 2016, 14 (1), e1002344. DOI: 10.1371/journal.pbio.1002344

(18) Wang, G.; de Jong, R. N.; van den Bremer, E. T.; Beurskens, F. J.; Labrijn, A. F.; Ugurlar, D.; Gros, P.; Schuurman, J.; Parren, P. W.; Heck, A. J. Molecular Basis of Assembly and Activation of Complement Component C1 in Complex with Immunoglobulin G1 and Antigen. Molecular Cell 2016, 63 (1), 135–145. DOI: 10.1016/j.molcel.2016.05.016

(19) Ugurlar, D.; Howes, S. C.; de Kreuk, B. J.; Koning, R. I.; de Jong, R. N.; Beurskens, F. J.; Schuurman, J.; Koster, A. J.; Sharp, T. H.; Parren, P.; et al. Structures of C1-IgG1 provide insights into how danger pattern recognition activates complement. Science 2018, 359 (6377), 794–797. DOI: 10.1126/science.aao4988

(20) Chromikova, V.; Mader, A.; Steinfellner, W.; Kunert, R. Evaluating the bottlenecks of recombinant IgM production in mammalian cells. Cytotechnology 2015, 67 (2), 343–356. DOI: 10.1007/s10616-014-9693-4

(21) Seeman, N. C. Nucleic Acid Junction and Lattices. Journal of Theoretical Biology 1982, 99, 237–247.

(22) Douglas, S. M.; Dietz, H.; Liedl, T.; Hogberg, B.; Graf, F.; Shih, W. M. Self-assembly of DNA into nanoscale three-dimensional shapes. Nature 2009, 459 (7245), 414–418. DOI: 10.1038/nature08016

(23) Wagenbauer, K. F.; Engelhardt, F. A. S.; Stahl, E.; Hechtl, V. K.; Stommer, P.; Seebacher, F.; Meregalli, L.; Ketterer, P.; Gerling, T.; Dietz, H. How We Make DNA Origami. Chembiochem 2017, 18 (19), 1873–1885. DOI: 10.1002/cbic.201700377

(24) Shaw, A.; Hoffecker, I. T.; Smyrlaki, I.; Rosa, J.; Grevys, A.; Bratlie, D.; Sandlie, I.; Michaelsen, T. E.; Andersen, J. T.; Hogberg, B. Binding to nanopatterned antigens is dominated by the spatial tolerance of antibodies. Nature Nanotechnology 2019, 14 (2), 184–190. DOI: 10.1038/s41565-018-0336-3

(25) Majumder, U.; Rangnekar, A.; Gothelf, K. V.; Reif, J. H.; LaBean, T. H. Design and construction of double-decker tile as a route to three-dimensional periodic assembly of DNA. Journal of the American Chemical Society 2011, 133 (11), 3843–3845. DOI: 10.1021/ja1108886

(26) He, Y.; Tian, Y.; Chen, Y.; Deng, Z.; Ribbe, A. E.; Mao, C. Sequence symmetry as a tool for designing DNA nanostructures. Angewandte Chemie International Edition 2005, 44 (41), 6694–6696. DOI: 10.1002/anie.200502193

(27) Zhang, F.; Jiang, S.; Wu, S.; Li, Y.; Mao, C.; Liu, Y.; Yan, H. Complex wireframe DNA origami nanostructures with multi-arm junction vertices. Nature Nanotechnology 2015, 10 (9), 779–784. DOI: 10.1038/nnano.2015.162

(28) Strasser, J.; de Jong, R. N.; Beurskens, F. J.; Wang, G.; Heck, A. J. R.; Schuurman, J.; Parren, P.; Hinterdorfer, P.; Preiner, J. Unraveling the Macromolecular Pathways of IgG Oligomerization and Complement Activation on Antigenic Surfaces. Nano Letters 2019, 19 (7), 4787–4796. DOI: 10.1021/acs.nanolett.9b02220

(29) Sharp, T. H.; Koster, A. J.; Gros, P. Heterogeneous MAC Initiator and Pore Structures in a Lipid Bilayer by Phase-Plate Cryo-electron Tomography. Cell Reports 2016, 15 (1), 1–8. DOI: 10.1016/j.celrep.2016.03.002

(30) Lubbers, R.; Oostindie, S. C.; Dijkstra, D. J.; Parren, P.; Verheul, M. K.; Abendstein, L.; Sharp, T. H.; de Ru, A.; Janssen, G. M. C.; van Veelen, P. A.; et al. Carbamylation reduces the capacity of IgG for hexamerization and complement activation. Clinical and Experimental Immunology 2020, 200 (1), 1–11. DOI: 10.1111/cei.13411

(31) Abendstein, L.; Dijkstra, D. J.; Tjokrodirijo, R. T. N.; van Veelen, P. A.; Trouw, L. A.; Hensbergen, P. J.; Sharp, T. H. Complement is activated by elevated IgG3 hexameric platforms and deposits C4b onto distinct antibody domains. Nature Communications 2023, accepted.

(32) Goldberg, B. S.; Ackerman, M. E. Antibody‐mediated complement activation in pathology and protection. Immunology & Cell Biology 2020, 98 (4), 305–317. DOI: 10.1111/imcb.12324

(33) Beum, P. V.; Lindorfer, M. A.; Beurskens, F.; Stukenberg, P. T.; Lokhorst, H. M.; Pawluczkowycz, A. W.; Parren, P. W.; van de Winkel, J. G.; Taylor, R. P. Complement activation on B lymphocytes opsonized with rituximab or ofatumumab produces substantial changes in membrane structure preceding cell lysis. The Journal of Immunology 2008, 181 (1), 822–832. DOI: 10.4049/jimmunol.181.1.822

(34) Gershov, D.; Kim, S.; Brot, N.; Elkon, K. B. C-Reactive protein binds to apoptotic cells, protects the cells from assembly of the terminal complement components, and sustains an antiinflammatory innate immune response: implications for systemic autoimmunity. Journal of Experimental Medicine 2000, 192 (9), 1353–1364. DOI: 10.1084/jem.192.9.1353

(35) Merle, N. S.; Noe, R.; Halbwachs-Mecarelli, L.; Fremeaux-Bacchi, V.; Roumenina, L. T. Complement System Part II: Role in Immunity. Frontiers in Immunology 2015, 6, 257. DOI: 10.3389/fimmu.2015.00257

(36) Pickering, M. C.; Botto, M.; Taylor, P. R.; Lachmann, P. J.; Walport, M. J. Systemic lupus erythematosus, complement deficiency, and apoptosis. Advances in Immunology 2000, 76, 227–324. DOI: 10.1016/s0065-2776(01)76021-x

(37) Mao, M.; Lin, Z.; Chen, L.; Zou, Z.; Zhang, J.; Dou, Q.; Wu, J.; Chen, J.; Wu, M.; Niu, L.; et al. Modular DNA-Origami-Based Nanoarrays Enhance Cell Binding Affinity through the “Lock-and-Key” Interaction. Journal of the American Chemical Society 2023, 145 (9), 5447–5455. DOI: 10.1021/jacs.2c13825

(38) Hellmeier, J.; Platzer, R.; Eklund, A. S.; Schlichthaerle, T.; Karner, A.; Motsch, V.; Schneider, M. C.; Kurz, E.; Bamieh, V.; Brameshuber, M.; et al. DNA origami demonstrate the unique stimulatory power of single pMHCs as T cell antigens. Proceedings of the National Academy of Sciences of the United States of America 2021, 118 (4). DOI: 10.1073/pnas.2016857118

(39) Wamhoff, E. C.; Knappe, G. A.; Burds, A. A.; Du, R. R.; Neun, B. W.; Difilippantonio, S.; Sanders, C.; Edmondson, E. F.; Matta, J. L.; Dobrovolskaia, M. A.; et al. Evaluation of Nonmodified Wireframe DNA Origami for Acute Toxicity and Biodistribution in Mice. ACS Applied Bio Materials 2023, 6 (5), 1960–1969. DOI: 10.1021/acsabm.3c00155

(40) Chandrasekaran, A. R. Nuclease resistance of DNA nanostructures. Nature Reviews Chemistry 2021, 5 (4), 225–239. DOI: 10.1038/s41570-021-00251-y

