## Supplemental Information for "DNA nanostructure-templated antibody complexes provide insights into the geometric requirements of human complement cascade activation"

### Experimental Section

#### DDT design and folding

DNA nanostructures were manually designed using the double-decker tile (DDT) first published by Majumder et al. as inspiration.[1] Molecular models of each design were created using the software program UCSF Chimera (version 1.16)[2] and UCSF ChimeraX (version 1.5)[3]. Each arm of the DDTs was modelled as four continuous dsDNA helices, placed 2 nm apart on a 2 × 2 grid, before being split at the locations of immobile Holliday junctions or termini. Holliday junctions were then formed using Isolde (version 1.0b3 within ChimeraX version 1.5)[4] and briefly simulated to relax the structures. The core $S_1$ and $S_2$ strands (or $S_3$ and $S_4$ in the case of DDT1; **Table S1**) were then linked together with the calculated number of thymine (T) bases,[5] before being allowed to relax in a short simulation within Isolde to yield the final molecular models.

DNA strands were ordered from Integrated DNA Technologies (Integrated DNA Technologies, BVBA, Belgium) with the sequences shown in **Table S1**, including purification using HPLC for strands up to 90 nucleotides (nt) and PAGE for longer strands. Antigen incorporation into the DDTs was performed during DNA synthesis of strand $S_5$, which included an internal DNP-TEG (2,4-dinitrophenyl-triethyleneglycol) modification. Similarly, cholesterol was incorporated into strand $S_8$ by extension of the 3' terminus with a TT linker and Cholesterol-TEG. The DDTs were produced by mixing 0.2 µM core strands ($S_1$ and $S_2$) with the correct molar ratios of the staple strands $S_{3\text{-}7}$ (without the $S_8$ strand); 1:1 (DDT1, 0.2 µM), 1:2 (DDT2, 0.4 µM), 1:3 (DDT3, 0.6 µM), 1:4 (DDT4, 0.8 µM), 1:5 (DDT5, 1 µM) and 1:6 (DDT6, 1.2 µM) in annealing buffer (20 mM Tris, 25 mM $MgCl_2$, 5 mM NaCl, pH 8.1). DDTs were folded using thermal annealing by gradually cooling down from 80°C to 4°C at a rate of 0.1°C every 1.3 min over a period of 17 h in a Bio-Rad C1000 Touch™ Thermal Cycler, and stored at 4°C. The $S_8$ strand, either only DNA or with the relevant cholesterol modification, was added 5 min before use to minimize aggregation caused by the cholesterol. The concentration of DNA nanostructures was measured using a NanoDrop™ 2000c spectrophotometer (Thermo Fisher Scientific, Waltham, Massachusetts, USA). DNA structures containing DNP were protected from light using aluminum foil.

#### DNA-DNP coupling and purification

As a positive control, the DNP antigen was directly linked to a separate $S_8$- strand containing a 5' amino group incorporated during synthesis. Amine-$S_8$-Cholesterol DNA strands (25 µM final concentration) and 1-(2,4-Dinitrophenylamino)-3,6,9,12 tetraoxapentadecanoic acid succinimidyl ester (DNP-$PEG_4$-NHS ester; 2.5 mM final concentration; Iris Biotech, Marktredwitz, Germany) were added to 1 × Phosphate-buffered saline (PBS) and dimethyl sulfoxide (DMSO; 1:1 ratio) and incubated overnight with agitation at 1,050 rpm in a ThermoMixer F2.0 (Eppendorf, Hamburg, Germany). For purification and buffer exchange to 1 ×PBS of DNP-$PEG_4$-$S_8$-Cholesterol, 10 kDa Amicon centrifugal filters were used. DNA concentration was determined using a NanoDrop™ 2000c spectrophotometer (Thermo Fisher Scientific, Waltham, Massachusetts, USA).

#### Antibody production

Recombinant anti-DNP immunoglobulin G (IgG) subclass 1 antibodies (Abs) were produced as previously described.[6] Briefly, sequences of IgG1 Abs against the hapten dinitrophenol (DNP) are based on the human constant domain of IgG1 (G1m(f) isotype) and the variable domain of monoclonal Ab G2a2.[7] Genes for heavy and κ-light chains were ordered from GeneArt (Thermo Fisher Scientific, Waltham, Massachusetts, USA) and cloned into a pcDNA3.3 vector (Thermo Fisher Scientific, Waltham, Massachusetts, USA) containing a signal peptide for secretion. Expi293F™ cells were transfected with 12.5 µg of the heavy chain plasmid and 12.5 µg of the light chain plasmid using the ExpiFectamine™ 293 Transfection Kit (Thermo Fisher Scientific, Waltham, Massachusetts, USA). Cells were incubated for 5 to 7 days, before centrifuging the transfection broth, followed by harvesting the supernatant and successively filtering using 0.45 µm and 0.2 µm filters (Cytiva, Marlborough, Massachusetts, USA). Abs were purified using a protein A column (Genscript, Piscataway, New Jersey, USA) and further concentrated and buffer-exchanged into 1 × PBS using 30 kDa Amicon centrifugal filters (Merck Millipore Sigma, Burlington, Massachusetts, USA). After purification, a NanoDrop™ 2000c spectrophotometer (Thermo Fisher Scientific, Waltham, Massachusetts, USA) was used to determine the concentration.

#### Characterization of DNA nanostructures and antibody complexes

To assess DNA nanostructure folding, 2% w/v UltraPure Agarose (Invitrogen, Thermo Fisher Scientific, Waltham, Massachusetts, USA) was dissolved in 1 × Tris-acetate-EDTA (TAE) buffer supplemented with 12 mM $MgCl_2$ (Sigma-Aldrich, St Louis, MO, USA). Strand $S_8$ was added in the correct molecular ratio and incubated with the pre-folded DNA nanostructures for 5 min at room temperature (10 ng/µl = 107.1 nM DDT1, 53.3 nM DDT2, 35.0 nM DDT3, 26.1 nM DDT4, 20.8 nM DDT5 and 17.1 nM DDT6 final concentration). Afterwards, 6 × loading dye (Thermo Fisher Scientific, Waltham, Massachusetts, USA) was combined with the mixture and loaded onto the 2% agarose (UltraPure Agarose, Invitrogen, Thermo Fisher Scientific, Waltham, Massachusetts, USA) gels. To check correctly incorporated DNP, anti-DNP IgG Abs (80 ng/µl = 533 nM final concentration) were added to the DNA nanostructures and incubated at room temperature for 15 min, before loading onto gels. Gels were run either on a Mini-PROTEAN Tetra Cell electrophoresis device (Bio-Rad Laboratories B.V., California, USA) at a constant voltage of 100 V for 1 h at 4°C or on a Sub-Cell GT Cell electrophoresis device (Bio-Rad Laboratories B.V., California, USA) at a constant voltage of 80 V for 2.5 h at 4°C. Agarose gels were stained for 10 min with GelRed® (Biotium, Inc., Fremont, California, USA) and imaged on a Gel Doc XR+ system (Bio-Rad Laboratories B.V., California, USA).For characterization of Ab binding using size exclusion chromatography (SEC), DNA nanostructures (100 nM DDT1, 50 nM DDT2, 33.3 nM DDT3, 25 nM DDT4, 20 nM DDT5 and 16.6 nM DDT6 final concentration), previously incubated with the strand $S_8$ for 5 min at room temperature and folded either ± DNP, were either loaded on an Increase 3.2/300 column (Cytiva, Marlborough, Massachusetts, USA) directly or first incubated at room temperature with anti-DNP IgG Abs (650 nM final concentration) for 20 min. The SEC column was equilibrated with

1 × PBS supplemented with 12 mM $MgCl_2$ on an Äkta™ pure system (Cytiva, Marlborough, Massachusetts, USA). Wavelengths 230 nm, 260 nm and 280 nm were detected simultaneously and the flow rate was set to 0.08 ml/min.

#### Liposome-based complement activity assay

All lipids were ordered from Avanti Polar Lipids (Alabama, USA). Liposomes which were used for DNA nanostructure binding contain no antigens, whereas liposomes only binding IgG Abs contained 1% 2,4 dinitrophenyl (DNP) as antigenic source. These were used for comparison in the cryo electron tomography setup (described below). Liposomes were produced as previously described.[6, 8] Briefly, lipid films composed of dimyristoylphosphatidylcholine (DMPC), dimyristoylphosphatidylglycerol (DMPG) and cholesterol (45:5:50 mol%) and DMPC, DMPG, cholesterol and dinitrophenyl-cap-dipalmitoylphosphatidylethanolamine (DNP-cap-PE) (44:5:50:1 mol%) were produced and, after incubating overnight on under a constant $N_2$ flow, rehydrated in 20 mM Sulforhodamine B (SRB; Sigma-Aldrich, St Louis, MO, USA) in 1 × PBS for 2 h at 37°C. Liposomes were generated by sonicating for 5 min at 60°C. Liposomes with encapsulated Sulforhodamine B were purified from free dye using Cytiva NAP™, NAP-25 columns (Cytiva, Marlborough, Massachusetts, USA). Rehydrated and purified liposomes were covered in aluminum foil and stored for a maximum of 3 weeks in the fridge until usage.

To measure DNA nanostructure-mediated complement activation, liposomes with encapsulated SRB were incubated with DNA nanostructures (10 nM, 25 nM, 50 nM, 75 nM and 100 nM final DNP concentration) for 10 min at room temperature. DNA nanostructures used in this assay were folded ± DNP and incubated with the $S_8$-Cholesterol strand for 5 min at room temperature before usage. DNA nanostructure concentration was normalized to DDT1 (i.e., 100 nM DDT1, 50 nM DDT2, etc.), which displays the DNP antigen. Samples were normalized to DDT1 to be able to compare Abs binding to the same amount of antigens. Next, 350 nM anti-DNP IgG was added and fluorescence intensity was measured for 100 s at 21°C. Finally, ice-cold normal human serum (NHS, Complement Technologies, Tyler, TX, USA; 10 % v/v final concentration) was added and fluorescence was monitored for 25 min at 21°C. For baseline correction, the fluorescent intensity of the first time point after NHS addition was used as a baseline and subtracted from all data points. Each sample was treated independently. Statistical analysis was performed in GraphPad Prism (version 9.3.1).

#### Western blot to detect C4 cleavage

DDT1-6 ± DNP were incubated at room temperature for 5 min with the $S_8$-Cholesterol DNA strand at the correct molecular ratio. Afterwards, the concentrations of DNA nanostructure were normalized to 100 nM of the antigen presenting strand $S_5$ (100 nM final DNP concentration) as before, by measuring the DNA concentration on a NanoDrop™ 200c spectrophotometer. Then, DDT1-6-Chol ± DNP were mixed with liposomes and incubated for 10 min at room temperature. The same antigen concentration was chosen for all experiments to be able to compare the same amount of nanopatterned Abs bound to DNA platforms. Next, 650 nM anti-DNP IgG Abs were added and incubated for 15 min at room temperature, followed by cooling to 4°C for 15 min. Ice-cold NHS (1% v/v final concentration) as a source for complement proteins was added and incubated for 30 min at 4°C. Liposomes, and any associated molecules, were purified via centrifugation using a Sigma 2-16k centrifuge (Sigma Laborzentrifugen GmbH, Harz, Germany) cooled to 4°C. Samples were centrifuged for 15 min at 20,000 g two times. Supernatants were removed after each centrifugation step and pellets were dissolved in ice-cold 1 × PBS. After the final washing step, samples were kept on ice until they were mixed with 2 × Laemmli buffer (Bio-Rad Laboratories B.V., California, USA) and dithiothreitol (0.1 M final concentration; DTT; Thermo Fisher Scientific, Waltham, Massachusetts, USA) and heated to 99°C for 10 min. This leads to denaturation of proteins and bursts the liposomes. This allows samples to enter the gel and makes them comparable to isolated, denatured proteins. Next, mixtures were loaded at equivalent normalized DNA concentrations (based on the concentration of DDT1, as above) concentration onto a 4 to 12% pre-casted Bis-Tris Protein Gels (Bolt™, Invitrogen™, Thermo Fischer Scientific Waltham, Massachusetts, USA). Gel electrophoresis was performed in 1× MOPS SDS buffer (51 mM 4-Morpholinepropanesulfonic acid, 3-(N-Morpholino) propanesulfonic acid, 50 mM Tris base, 3.45 mM sodium dodecyl sulfate, 1.05 mM EDTA; pH 7.7) in an Invitrogen™ Mini Gel Tank electrophoresis device (Thermo Fisher Scientific, Waltham, Massachusetts, USA) for 35 min at 200 V at room temperature.

After gel electrophoresis, gels were either stained using silver-stain (Pierce™ Silver Stain Kit Thermo Fisher Scientific, Waltham, Massachusetts, USA) or proteins were transferred to a Hybond C blotting membrane (Amersham Life Sciences, Buckinghamshire, United Kingdom) for 3 h in an ice bath in blotting buffer (95.9 mM Glycine, 12 mM Tris and 2.5 M Methanol) at 0.35 A per blot on an Invitrogen™ Mini Gel Tank electrophoresis device (Thermo Fisher Scientific, Waltham, Massachusetts, USA). Then, blots were washed three times for 5 min using washing buffer (1 × PBS plus 0.1% Tween-20 w/v) before blocking in blocking buffer (1 × PBS plus 0.1% Tween-20 w/v and 5% milk w/v) for 30 min at room temperature. Afterwards, C4 complement protein was detected by incubating the blots with anti-human C4 goat Abs (1:128,000 diluted in blocking buffer, Complement Technology, Tyler, TX, USA) overnight on a vertical tube shaker at 4°C. Blots were washed three times for 5 min using the blocking buffer. To visualize the C4 Abs, an anti-goat donkey secondary horse radish peroxidase (HRP) Abs (1:1,000 diluted in blocking buffer; Invitrogen, Thermo Fisher Scientific, Waltham, Massachusetts, USA) was added and incubated at room temperature on a vertical tube roller for 30 min. Next, blots were washed two times for 5 min in block buffer, followed by four times washing for 5 min in washing buffer. Finally, blots were treated for 5 min with 2 ml per blot Clarity Western ECL Substrate (Bio-Rad Laboratories B.V., California, USA) before imaging on a ChemiDoc MP (Bio-Rad Laboratories B.V., California, USA).

After visualizing the C4 complement protein, detection Abs were stripped from the blots. Blots were incubated twice with stripping buffer (199.8 mM Glycine, 3.5 mM SDS, 8.1 mM Tween-20, pH 2.2) for 15 min then twice washed with 1 × PBS for 10 min, followed by three times washing with washing buffer for 5 min and blocking in blocking buffer for 30 min at room temperature. Next, C1q complement protein was detected using anti-human C1q rabbit Abs (1:1,000 diluted in blocking buffer, Dako, Denmark) at room temperature on a vertical tube roller for 30 min, followed by three times washing in block buffer and 30 min incubation of anti-rabbit goat secondary HRP Abs (1:1,000 diluted in blocking buffer; Dako, Denmark) on a vertical roller at room temperature. Afterwards, blots were again washed two times for 5 min in the blocking buffer and then four times for 5 min in the wash buffer, before developing the blot using the Clarity Western ECL Substrate. To visualize IgG Abs the blot was stripped as described above, followed by incubating the blots with anti-human IgG (H+L) goat Abs (1:1,000 diluted in blocking buffer, Thermo Fisher Scientific, Waltham, Massachusetts, USA) overnight at 4°C on a vertical tuber roller. Blots were washed three times for 5 min using blocking buffer, followed by incubating the blots with anti-goat donkey secondary HRP Abs (1:1,000 diluted in blocking buffer) for 30 min at room temperature before developing as described above.

#### Negative stain electron microscopy and image analysis

Samples, 10 µl, were loaded on freshly plasma-cleaned 200 mesh carbon-coated copper grids (Electron Microscopy Sciences, PA, USA), incubated for 1 min and blotted using Whatman paper (Sigma-Aldrich, St Louis, MO, USA), before washing with water twice. Samples were stained using 3 µl 2 % uranyl formate (Spi® Supplies West Chester, Pennsylvania, USA) for 20 s, after which excess uranyl formate was blotted away. Negative stain micrographs were collected on a FEI Tecnai T12 BioTwin with $LaB_6$ electron source, operating at 120 kV on a FEI Eagle 4k × 4k CCD camera at 49,000 × magnification, with a nominal defocus of -1.5 µm and a pixel size of 4.546 Å. EMAN2 (version 2.91)[9] was used for particle picking and to generate class averages.

#### Cryo-electron microscopy sample preparation

The $S_8$-Cholesterol strand, at a final concentration of 1.2 µM, was added to 0.2 µM DDT6 and incubated for 5 min at room temperature. Then, 50 nM DNA nanostructures were added to liposomes and incubated for 10 min at room temperature. Afterwards, anti-DNP IgG Abs (350 nM final concentration) were added and incubated for 15 min at room temperature, before cooling down to 4°C for 15 min. Next, ice-cold 1 × PBS or NHS (1.5 % v/v final concentration) was added and incubated at 4°C for 30 min. Just prior to vitrification, 5 nm bovine serum albumin-coated gold colloids (Department of Cell Biology, UMC, Utrecht, the Netherlands) were added. Grids were loaded into a Leica EMGP (Leica Microsystems, Wetzlar, Germany) and 6 µl of the sample were directly applied onto freshly plasma-cleaned 200 mesh copper grids with lacey-carbon support (Electron Microscopy Sciences, PA, USA). Samples were incubated for 60 s at 4°C at 65% humidity, and blotted for 1.5 s from the back before vitrification in liquid ethane. Grids were clipped and stored in liquid nitrogen until usage.

#### Data collection and tomogram reconstruction of cryo-electron microscopy samples

Tilt series were acquired on either a Talos Arctica (Thermo Fisher Scientific; at Leiden University Medical Centre) operated at 200 kV with a K3 direct electron detector operating in counting mode and a Bioquantum energy filter (Gatan) set to a slit width of 20 eV or a Titan Krios G1 (Thermo Fisher Scientific; at Netherlands Centre for Electron Nanoscopy) operated at 300 kV with a FEI Falcon 3 direct electron detector, both operated in counting mode. For data acquisition at the Talos Arctica, the Tomo 5.5.0 data acquisition software package (Thermo Fisher Scientific) was used. Tilt series were collected at 49,000 × magnification with a pixel size of 1.74 Å using a dose symmetric scheme (± 57°, in 3° increments). The nominal defocus was set between -4 to -6 µm and tracking was performed before each tilt acquisition. A total dose of 60 $e^-/Å^2$ was used for each tilt series. For data acquisition at the Titan Krios, Tomography 4.0.0 software (Thermo Fisher Scientific) was used at 22,500 × magnification with a pixel size of 3.0975 Å using a dose symmetric scheme (± 54°, in 3° increments). The nominal defocus was acquired between -4 to -6 µm and tracking was performed before each tilt acquisition. A total dose of 60 $e^-/Å^2$ was used for each tilt series. For tomogram reconstruction, IMOD (version4.11) [10] was used, and EMAN2 (version 2.91)[9] e2spt_boxer_old.py was used to visualize tomograms.

### Supplementary Figures

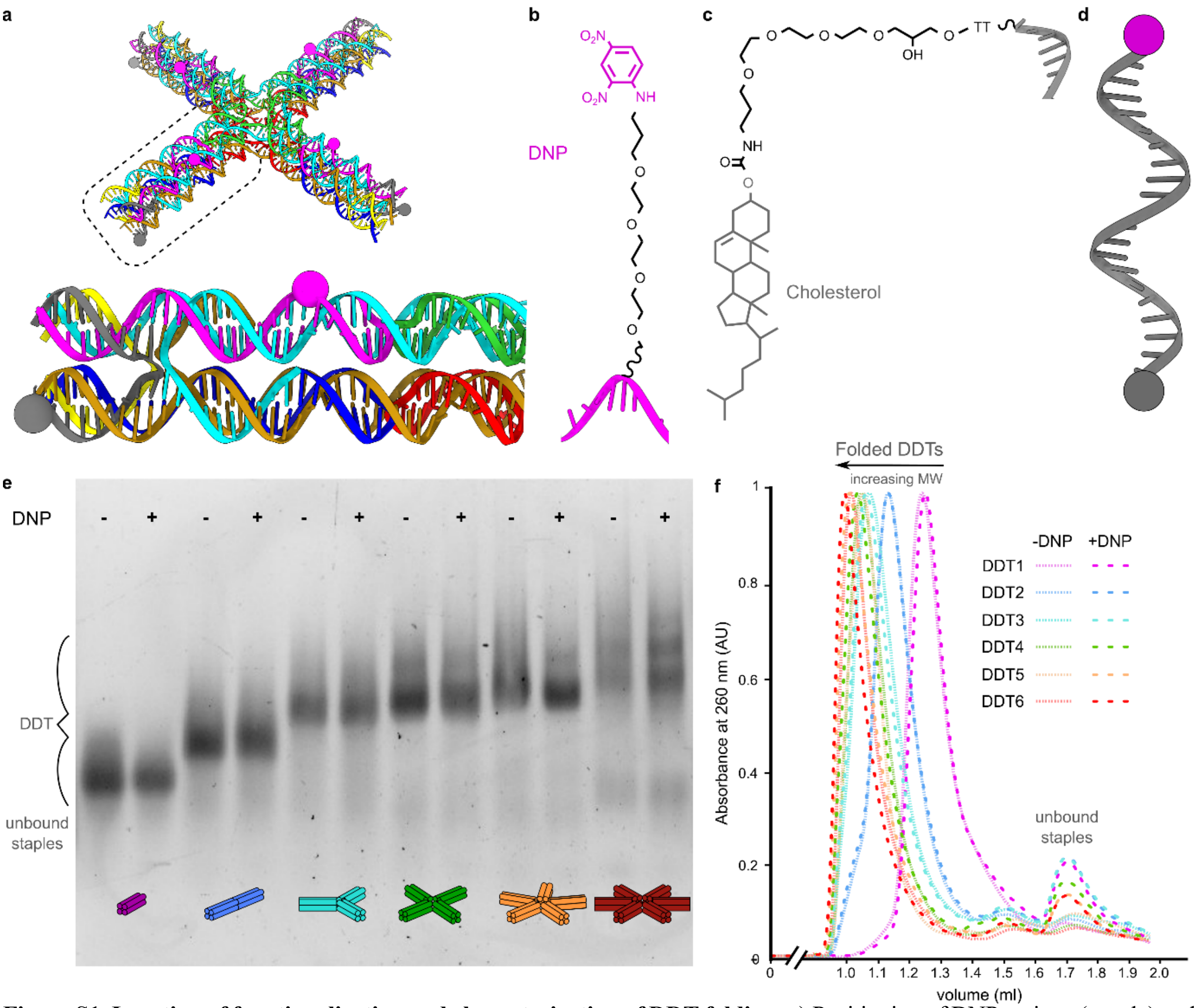


**Figure S1. Location of functionalization and characterization of DDT folding. a**) Positioning of DNP antigen (purple) and cholesterol (grey) functionalisations on ssDNA strands. **b & c**) chemical structures of DNP antigen and cholesterol modified strands. **d**) Schematic of single stranded positive control DNA strand functionalized with both DNP (purple) and cholesterol (grey). **e**) Agarose gel of folded DDT1-6 ± DNP functionalization. **f**) Normalized SEC trace at 260 nm of folded DDT1-6 ± DNP functionalization after folding, without previous purification.

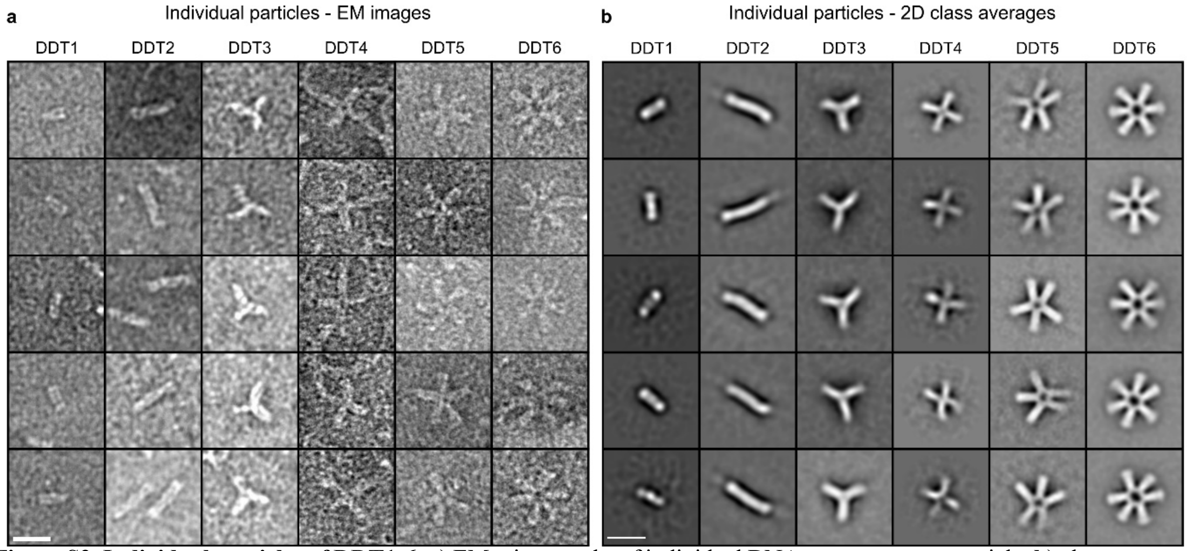


**Figure S2. Individual particles of DDT1-6. a**) EM micrographs of individual DNA nanostructures particle. **b**) class averages generated in EMAN2 of DNA nanostructures identified on EM images. Scale bar represents 20 nm for all images**.**

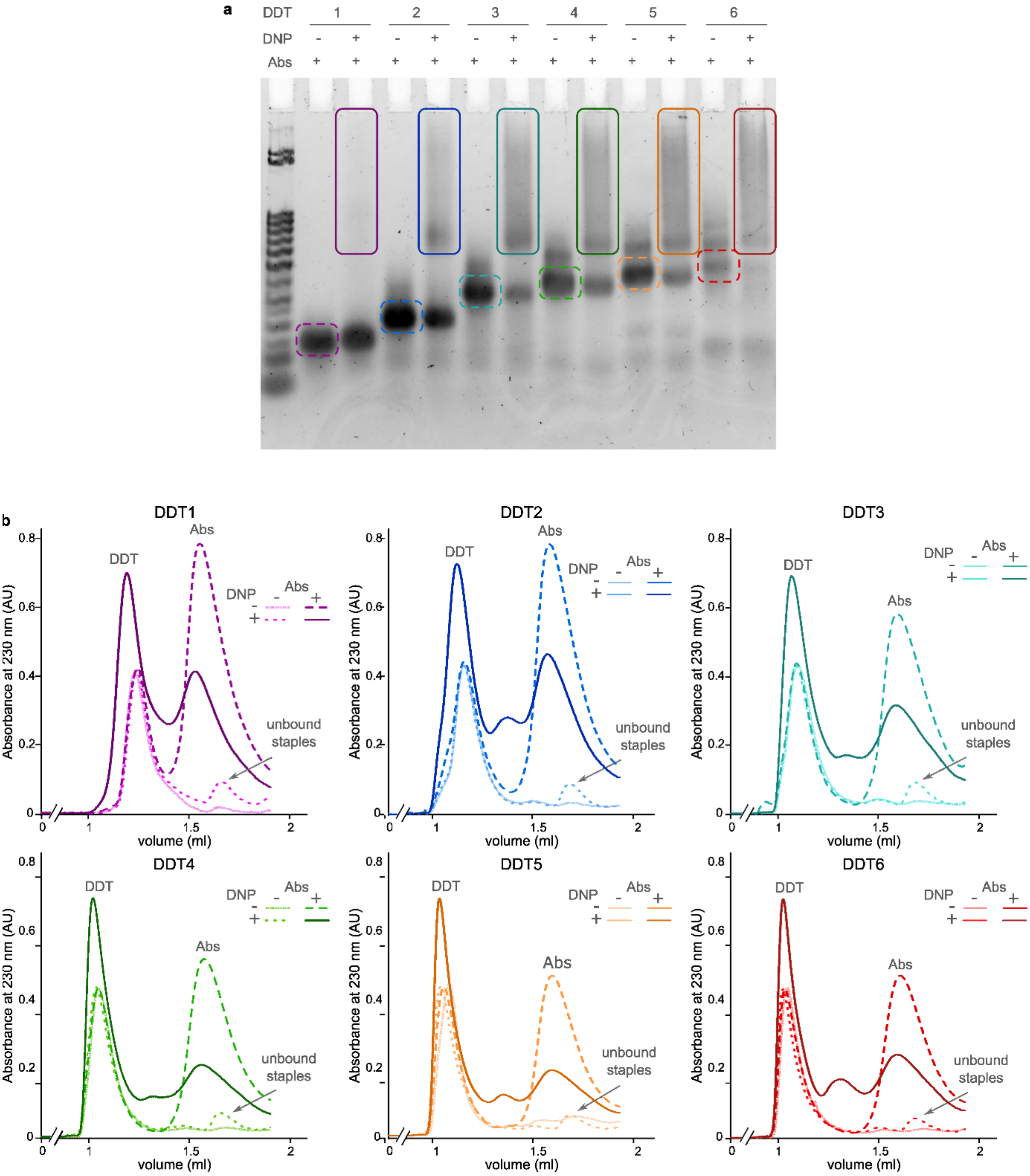


**Figure S3. Characterization of Abs binding to DDT nanostructures in the presence or absence of DNP antigen. a)** Agarose gel of folded DDT1-6 ± DNP functionalization preincubated with anti-DNP IgG Abs. **b**) Normalized SEC trace at 230 nm of folded DDT1-6 ± DNP functionalization, without previous purification incubated with anti-DNP IgG Abs.

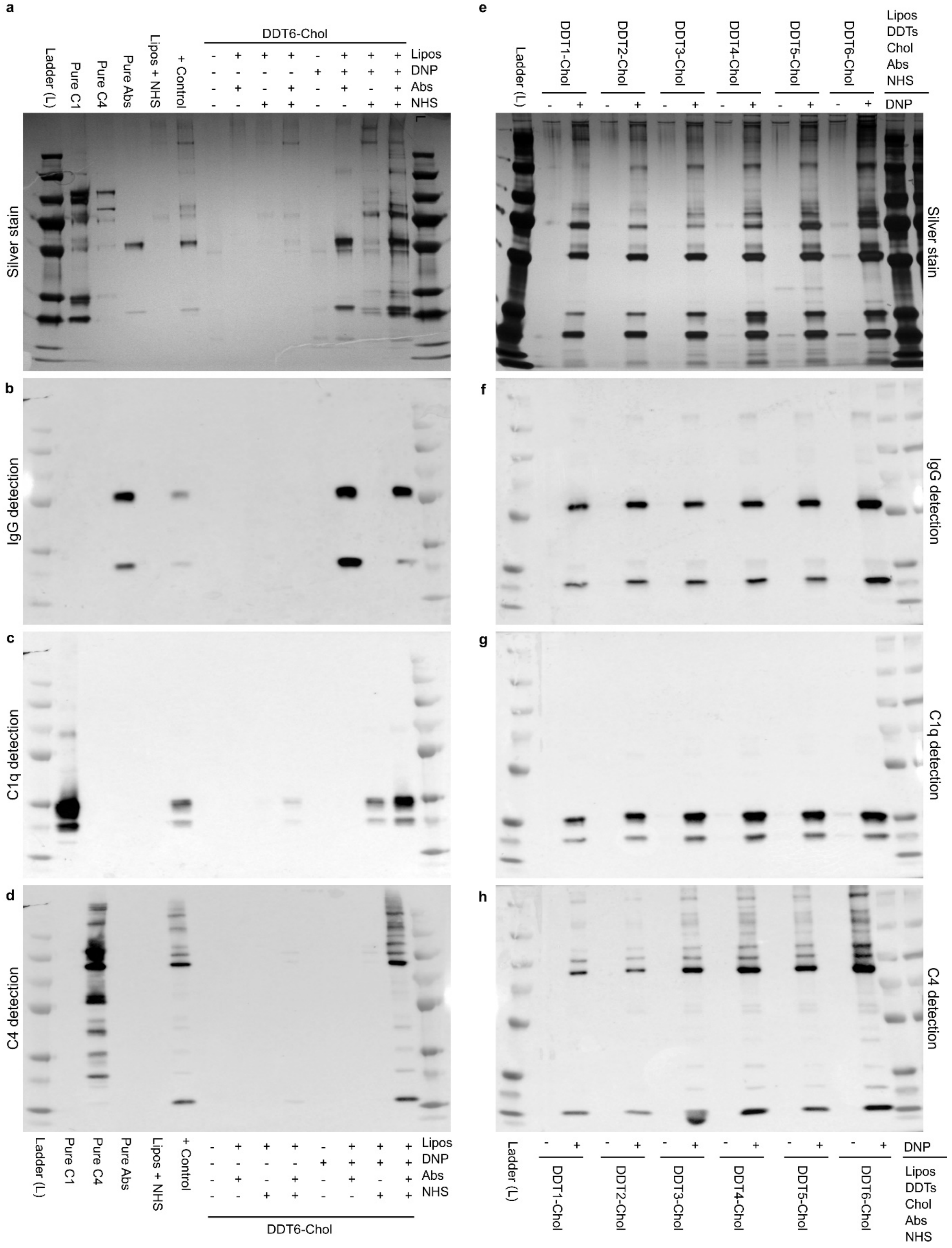


**Figure S4. DDT1-6 mediated classical complement activation measured via C4 cleavage. a**) Silver-stained PAGE of purified proteins on isolated liposomes after incubation with DDT6-Chol, folded ± DNP and the components shown. The first 5 lanes contain control proteins. **b-d**) Western blot detecting IgG **b**), C1q **c**) and C4 **d**) on purified proteins on liposomes after incubation with DDT6 containing cholesterol, folded ± DNP and NHS. The first 5 lanes contain control proteins. **e**) Silver-stained PAGE of purified proteins on isolated liposomes after incubation with DDT1-6-Cholesterol, folded ± DNP and the components shown. **f-h**) Western blot detecting IgG **f**), C1q **g**) and C4 **h**) on purified proteins on liposomes after incubation with DDT1-6-Chol + DNP and NHS. Sample compositions are described above and below the gels and blots, respectively.

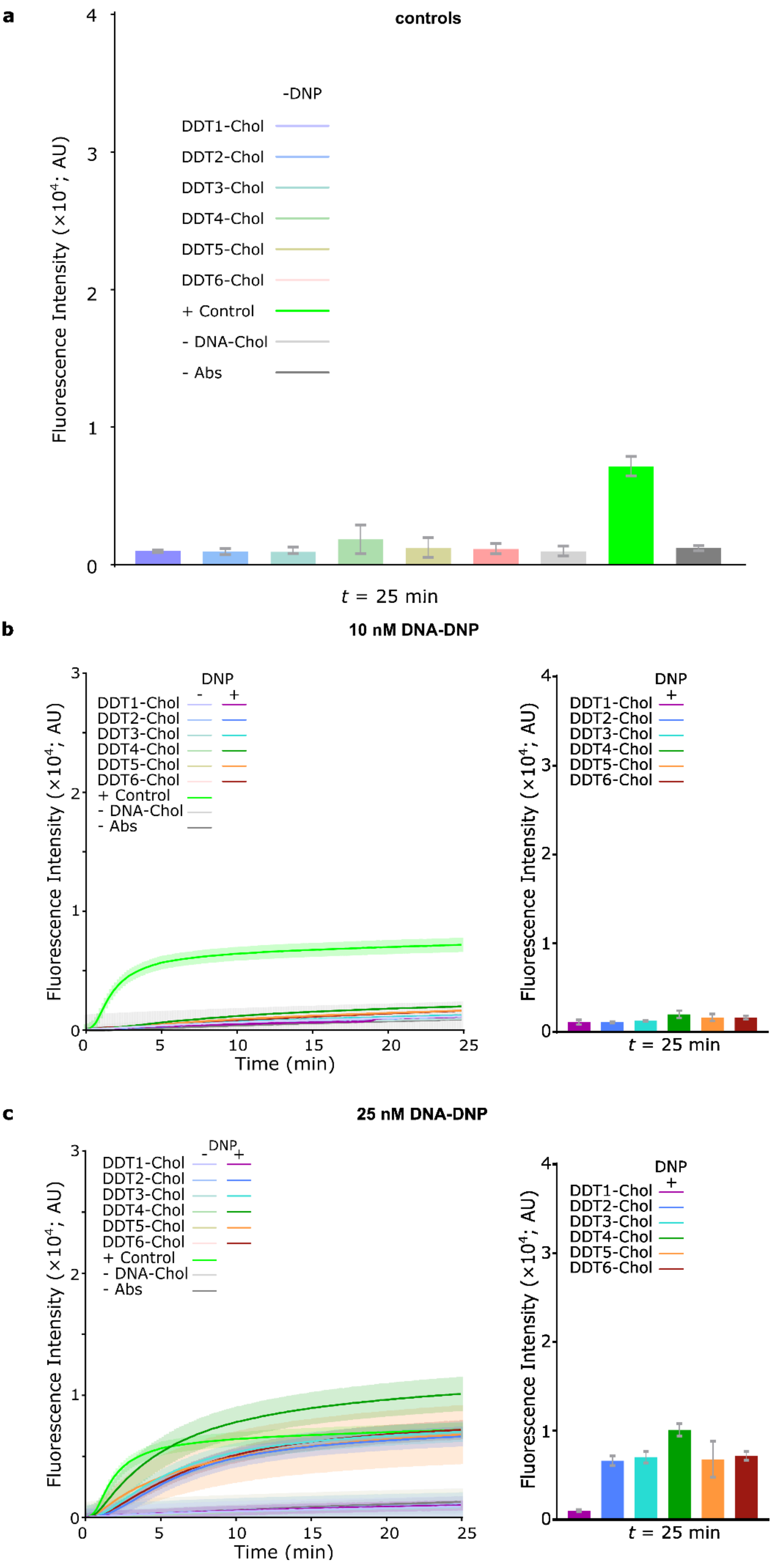
a
controls
-DNP
DDT1-Chol
DDT2-Chol
DDT3-Chol
DDT4-Chol
DDT5-Chol
DDT6-Chol
+ Control
- DNA-Chol
- Abs
Fluorescence Intensity (×10^4; AU)
t = 25 min
b
10 nM DNA-DNP
DNP
Time (min)
c
25 nM DNA-DNP

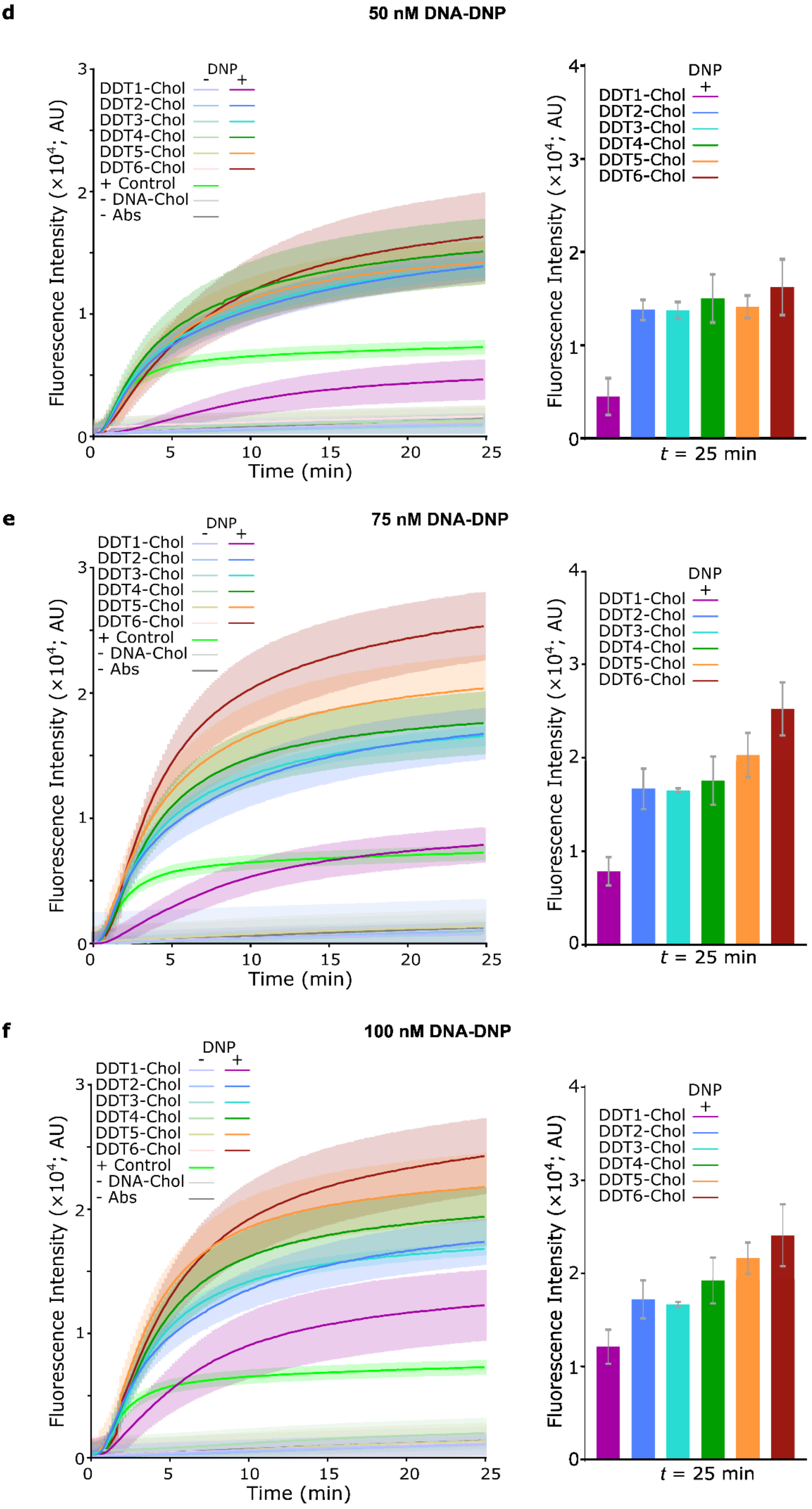
d
50 nM DNA-DNP
e
75 nM DNA-DNP
f
100 nM DNA-DNP
DNP
- +
DDT1-Chol
DDT2-Chol
DDT3-Chol
DDT4-Chol
DDT5-Chol
DDT6-Chol
+ Control
- DNA-Chol
- Abs
Fluorescence Intensity (×10^4; AU)
Time (min)
0 5 10 15 20 25
t = 25 min

**Figure S5. Assessing complement activation by DNA-templated antibody complexes using liposome lysis assays.** **a**) Graph representing all controls, after a 25 min incubation, used in this assay. **b**) Liposome lysis assay showing an increase in fluorescence caused by DNA-templated antibody binding at 10 nM DDT (left). Graphs representing the fluorescence intensity after 25 min of the assay (right). **c**) Liposome lysis assay showing an increase in fluorescence caused by DNA-templated antibody binding at 25 nM DDT (left). Graphs representing the fluorescence intensity after 25 min of the assay (right). **d**) Liposome lysis assay showing an increase in fluorescence caused by DNA-templated antibody binding at 50 nM DDT (left). Graphs representing the fluorescence intensity after 25 min of the assay (right). **e**) Liposome lysis assay showing an increase in fluorescence caused by DNA-templated antibody binding at 75 nM DDT (left). Graphs representing the fluorescence intensity after 25 min of the assay (right). **f**) Liposome lysis assay showing an increase in fluorescence caused by DNA-templated antibody binding at 100 nM DDT (left). Graphs representing the fluorescence intensity after 25 min of the assay (right). Error bars in all panels represent the standard error of at least 3 independent replicates. The positive control (+ *control*) as well as the negative controls (- *DNA-Chol* and – *Abs*) are the same for every graph.

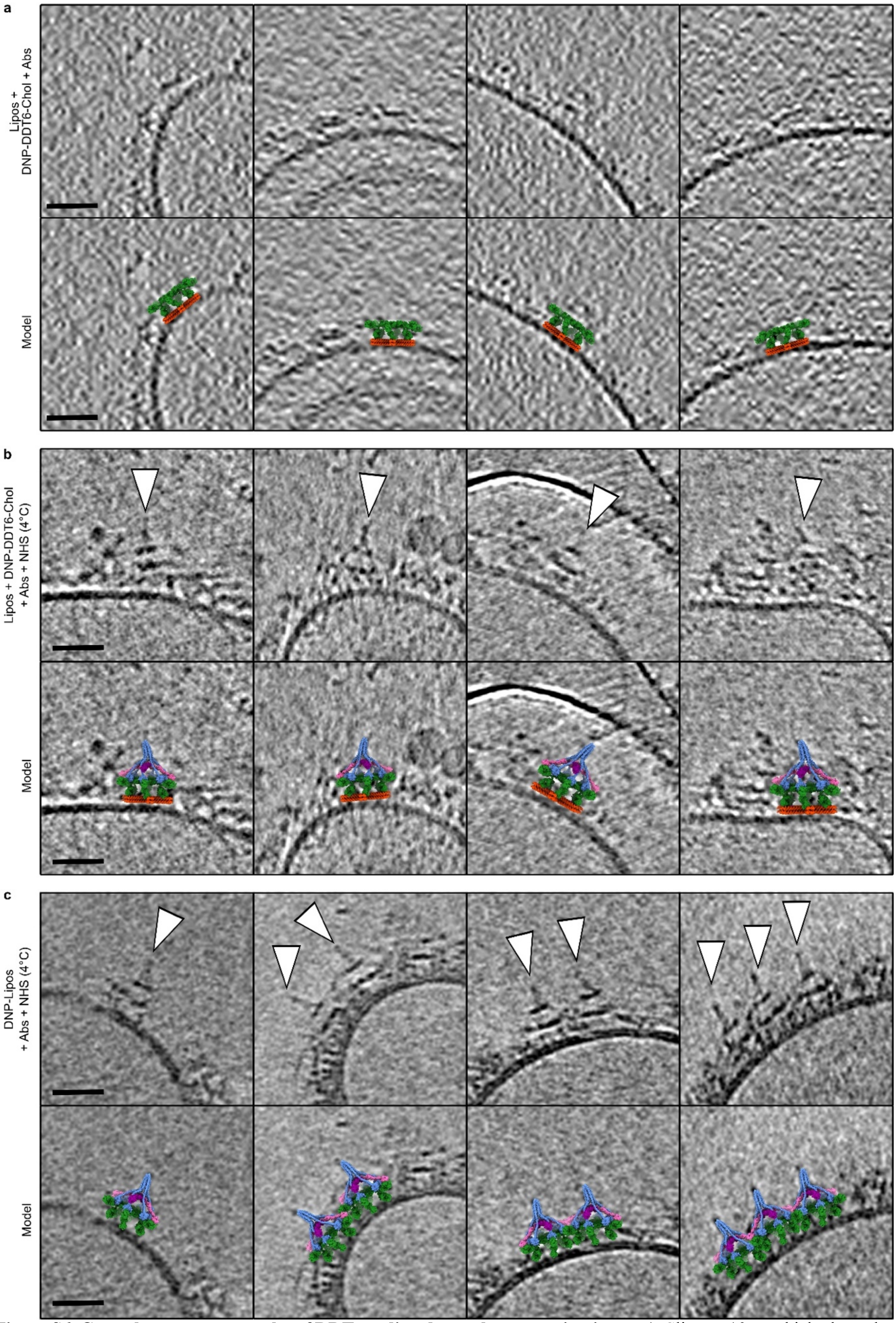


**Figure S6. Cryo-electron tomography of DDT-mediated complement activation. a-c**) Slices, ~10 nm thick, through cryo-electron tomograms (top) and overlaid model (bottom). **a**) Tomographic slices of DNP-DDT6-Chol with bound anti-DNP IgG Abs on the surface of liposomes. **b**) Tomographic slices of DNP-DDT6-Chol incubated with anti-DNP IgG Abs and NHS at 4 °C on the surface of liposomes. **c**) Tomographic slices of liposomes presenting DNP incubated with anti-DNP IgG Abs and NHS at 4 °C. All scale bars represent 30 nm.

### Supplementary Tables

**Table S1. DNA sequences used in this study for DNA nanostructures and control strands.**

| specification | | name | sequence | color |
|---|---|---|---|---|
| **Core strands** | **DDT1** | **$S_1$-mono** | AAC CAC TAG ATT GGC GAG ACG | green |
| | | **$S_2$-mono** | AGC ACC AAT CGT AGG CTT GCC | red |
| | **DDT2** | **$S_1$-bi** | AAC CAC TAG ATT GGC GAG ACG AAC CAC TAG ATT GGC GAG ACG | green |
| | | **$S_2$-bi** | AGC ACC AAT CGT AGG CTT GCC AGC ACC AAT CGT AGG CTT GCC | red |
| | **DDT3** | **$S_1$-tri_T3** | AAC CAC TAG ATT GGC TTT GAG ACG AAC CAC TAG ATT GGC TTT GAG ACG AAC CAC TAG ATT GGC TTT GAG ACG | green |
| | | **$S_2$-tri_T3** | AGC ACC AAT CGT AGG TTT CTT GCC AGC ACC AAT CGT AGG TTT CTT GCC AGC ACC AAT CGT AGG TTT CTT GCC | red |
| | **DDT4** | **$S_1$-tet_T4** | AAC CAC TAG ATT GGC TTTT GAG ACG AAC CAC TAG ATT GGC TTTT GAG ACG AAC CAC TAG ATT GGC TTTT GAG ACG AAC CAC TAG ATT GGC TTTT GAG ACG | green |
| | | **$S_2$-tet_T4** | AGC ACC AAT CGT AGG TTTT CTT GCC AGC ACC AAT CGT AGG TTTT CTT GCC AGC ACC AAT CGT AGG TTTT CTT GCC AGC ACC AAT CGT AGG TTTT CTT GCC | red |
| | **DDT5** | **$S_1$-pent_T5** | AAC CAC TAG ATT GGC TTTTT GAG ACG AAC CAC TAG ATT GGC TTTTT GAG ACG AAC CAC TAG ATT GGC TTTTT GAG ACG AAC CAC TAG ATT GGC TTTTT GAG ACG AAC CAC TAG ATT GGC TTTTT GAG ACG | green |
| | | **$S_2$-pent_T5** | AGC ACC AAT CGT AGG TTTTT CTT GCC AGC ACC AAT CGT AGG TTTTT CTT GCC AGC ACC AAT CGT AGG TTTTT CTT GCC AGC ACC AAT CGT AGG TTTTT CTT GCC AGC ACC AAT CGT AGG TTTTT CTT GCC | red |
| | **DDT6** | **$S_1$-hex_T7** | AAC CAC TAG ATT GGC TTTTTTT GAG ACG AAC CAC TAG ATT GGC TTTTTTT GAG ACG AAC CAC TAG ATT GGC TTTTTTT GAG ACG AAC CAC TAG ATT GGC TTTTTTT GAG ACG AAC CAC TAG ATT GGC TTTTTTT GAG ACG AAC CAC TAG ATT GGC TTTTTTT GAG ACG | green |
| | | **$S_2$-hex_T7** | AGC ACC AAT CGT AGG TTTTTTT CTT GCC AGC ACC AAT CGT AGG TTTTTTT CTT GCC AGC ACC AAT CGT AGG TTTTTTT CTT GCC AGC ACC AAT CGT AGG TTTTTTT CTT GCC AGC ACC AAT CGT AGG TTTTTTT CTT GCC AGC ACC AAT CGT AGG TTTTTTT CTT GCC | red |
| **Staple strands** | **general** | **$S_3$-blunt** | ACC GTC ACT TCG CTT CTG TAT TCG ATC CGG TTC GTC TCG CCA ATC TAG TAT CGC GCA GTC GGC ACG TGT AGT TGA C | cyan |
| | | **$S_4$** | ATG CTA AGG AAG TGA CGG TCA GAC ACC TGC TGG CAA GCC TAC GAT TGG ACG CGG TAT GGA ACG TAC ACG TGC CG | brown |
| | | **$S_5$-blunt** | ACT GCG CGA TGG ATC GAA TAC AGA AGC GTC ATC TCC G | pink |
| | | **$S_6$** | CTG GAG TAA GTA CGT TCC ATA CCG CGT GGT GTC TG | blue |
| | | **$S_7$-blunt** | GTC AAC TAC TTA CTC CAG | yellow |
| | | **$S_8$-blunt** | CGG AGA TGA CCT TAG CAT | black |
| **Modified strands** | **DDT1** | **$S_3$-blunt_mono** | ACC GTC ACT TCG CTT CTG TAT TCG ATC CGG TTC GTC TC TTTTTTTTT G CCA ATC TAG TAT CGC GCA GTC GGC ACG TGT AGT TGA C | cyan |
| | | **$S_4$_mono** | ATG CTA AGG AAG TGA CGG TCA GAC ACC TGC TGG CAA G TTTTTTTTT CC TAC GAT TGG ACG CGG TAT GGA ACG TAC ACG TGC CG | brown |
| | **general** | **$S_5$-DNP-N** | ACT GCG CGA TGG ATC GA/iDNPTEG/ TAC AGA AGC GTC ATC TCC G | pink |
| | | **$S_8$-Chol** | CGG AGA TGA CCT TAG CAT TT/3CholTEG/ | black |
| | **control** | **Amine-$S_8$-Chol** | /5AmMC6/CGG AGA TGA CCT TAG CAT TT/3CholTEG/ | black |

### References

(1) Majumder, U.; Rangnekar, A.; Gothelf, K. V.; Reif, J. H.; LaBean, T. H. Design and construction of double-decker tile as a route to three-dimensional periodic assembly of DNA. *Journal of the American Chemical Society* **2011**, *133* (11), 3843-3845. DOI: 10.1021/ja1108886

(2) Pettersen, E. F.; Goddard, T. D.; Huang, C. C.; Couch, G. S.; Greenblatt, D. M.; Meng, E. C.; Ferrin, T. E. UCSF Chimera--a visualization system for exploratory research and analysis. *Journal of Computational Chemistry* **2004**, *25* (13), 1605-1612. DOI: 10.1002/jcc.20084

(3) Pettersen, E. F.; Goddard, T. D.; Huang, C. C.; Meng, E. C.; Couch, G. S.; Croll, T. I.; Morris, J. H.; Ferrin, T. E. UCSF ChimeraX: Structure visualization for researchers, educators, and developers. *Protein Science* **2021**, *30* (1), 70-82. DOI: 10.1002/pro.3943

(4) Croll, T. I. ISOLDE: a physically realistic environment for model building into low-resolution electron-density maps. *Acta Crystallographica Section D* **2018**, *D74* (Pt 6), 519-530. DOI: 10.1107/S2059798318002425

(5) Zhang, F.; Jiang, S.; Wu, S.; Li, Y.; Mao, C.; Liu, Y.; Yan, H. Complex wireframe DNA origami nanostructures with multi-arm junction vertices. *Nature Nanotechnology* **2015**, *10* (9), 779-784. DOI: 10.1038/nnano.2015.162

(6) Abendstein, L.; Dijkstra, D. J.; Tjokrodirijo, R. T. N.; van Veelen, P. A.; Trouw, L. A.; Hensbergen, P. J.; Sharp, T. H. Complement is activated by elevated IgG3 hexameric platforms and deposits C4b onto distinct antibody domains. *Nature Communications* **2023**, *accepted*.

(7) Gonzalez, M. L.; Frank, M. B.; Ramsland, P. A.; Hanas, J. S.; Waxman, F. J. Structural analysis of IgG2A monoclonal antibodies in relation to complement deposition and renal immune complex deposition. *Molecular Immunology* **2003**, *40* (6), 307-317. DOI: 10.1016/s0161-5890(03)00167-6

(8) Lubbers, R.; Oostindie, S. C.; Dijkstra, D. J.; Parren, P.; Verheul, M. K.; Abendstein, L.; Sharp, T. H.; de Ru, A.; Janssen, G. M. C.; van Veelen, P. A.; et al. Carbamylation reduces the capacity of IgG for hexamerization and complement activation. *Clinical and Experimental Immunology* **2020**, *200* (1), 1-11. DOI: 10.1111/cei.13411

(9) Galaz-Montoya, J. G.; Flanagan, J.; Schmid, M. F.; Ludtke, S. J. Single particle tomography in EMAN2. *Journal of Structural Biology* **2015**, *190* (3), 279-290. DOI: 10.1016/j.jsb.2015.04.016

(10) Kremer, J. R.; Mastronarde, D. N.; McIntosh, J. R. Computer Visualization of Three-Dimensional Image Data Using IMOD. *Journal of Structural Biology* **1995**, *116*, 71-76.
